# Inhibiting retinoic acid mitigates vision loss in a mouse model of retinal degeneration

**DOI:** 10.1101/2021.08.09.455683

**Authors:** Michael Telias, Kevin Sit, Daniel Frozenfar, Benjamin Smith, Arjit Misra, Michael J Goard, Richard H Kramer

## Abstract

In degenerative retinal disorders, rod and cone photoreceptors die, causing vision impairment and blindness. Downstream neurons survive but undergo morphological and physiological remodeling, with some retinal ganglion cells (RGC) exhibiting heightened spontaneous firing. Retinoic acid (RA) has been implicated as the key signaling molecule that induces RGC hyperactivity, obscuring RGC light responses and reducing light avoidance behaviors triggered by residual rods and cones. However, evidence that RA-dependent remodeling corrupts image-forming vision has been lacking. Here we show that disulfiram, an FDA-approved drug that inhibits RA synthesis, and BMS 493, an RA receptor (RAR) inhibitor, reduce RGC hyperactivity and augment image detection in visually impaired mice. Functional imaging of visual cortical neurons shows that disulfiram and BMS 493 sharpen orientation-tuning and strengthen response fidelity to naturalistic scenes. These findings establish a causal link between RA-induced retinal hyperactivity and vision impairment and define molecular targets and candidate drugs for boosting image-forming vision in retinal degeneration.

## Introduction

Age-related macular degeneration (AMD) and retinitis pigmentosa (RP) are the most prevalent photoreceptor degenerative disorders, impairing vision or causing blindness in hundreds of millions of people worldwide^1, 2^. Both AMD and RP progress gradually, with retinal light responses and visual perception declining over years to decades. Despite the slow deterioration of rods and cones, most retinal ganglion cells (RGCs) survive and maintain synaptic connectivity with the brain until late stages in disease^3–5^. This has made RGCs a potential substrate for artificial vision restoration by optoelectronics^6, 7^, optogenetics^8, 9^, and optopharmacology^10, 11^. Because vision restoration technologies supplant light responses initiated by residual rods and cones, treatments based on these technologies are being deployed only in a small fraction of people with late-stage retinal degeneration. For patients whose vision is severely impaired but not yet eliminated, treatments for improving sight remain an unmet medical need.

Studies on mouse, rat, and rabbit models of RP show that while downstream retinal neurons do survive, they nonetheless undergo morphological and physiological remodeling. Months after photoreceptors are lost, new dendritic branches begin to sprout from downstream neurons and their cell body positions start to change, mirroring remodeling events that occur over years to decades in advanced human RP^12–15^. Much earlier, within days of photoreceptor loss, some RGCs become hyperactive, firing spontaneously at up to 8 times their normal rates in healthy retinas^16, 17^. RGC hyperactivity results partly from changes in presynaptic neurons^18, 19^ and partly from changes intrinsic to RGCs. These include up-regulation of voltage-gated ion channels, increasing membrane excitability^11^ and up-regulation of ligand-gated channels, increasing resting membrane permeability^20^. Recent studies indicate that retinoic acid (RA) is the signal that triggers morphological^21^ and physiological^17, 22^ remodeling. Adding exogenous RA to healthy retinas mimics remodeling, adding RA receptor (RAR) inhibitors to degenerated retinas suppresses remodeling, and increased RA-induced gene expression can be detected in degenerated retinas^17^. Local atrophy of photoreceptors, induced by subretinal implantation of a metallic chip, also caused elevation of RA-induced gene expression in corresponding RGCs, leading to local hyperactivity^23^. This suggests that RA-induced hyperactivity is a common sequel to photoreceptor loss, whether the underlying cause is hereditary, as in RP, or environmental, as in many cases of AMD.

The functional consequences of remodeling on information processing and visual perception have remained uncertain^24^. Heightened spontaneous RGC firing could mask responses triggered by residual photoreceptors, as has been observed in the rd10 mouse model of RP^17^. However, degeneration-induced hyperactivity^18^ and hyperpermeability^20^ apply only to Off-RGCs, and not other RGC types. Heightened spontaneous activity in Off-RGCs is predicted to compress the dynamic range to light decrements that normally provoke firing and expand the dynamic range to light increments that normally suppress firing. But how these seemingly opposite effects might impact higher order visual information processing in the brain is unknown. The effects of hyperactivity on visual perception are also unclear. Genetically inhibiting RAR increased light-avoidance behavior in vision-impaired rd10 mice^17^, but whether this reflects improved visual acuity or enhanced photophobia has been unclear.

There is direct and circumstantial evidence that morphological and physiological remodeling, including retinal hyperactivity, applies to humans with degenerative blindness, similar to animal models^14, 24^. RP patients report persistent photopsias^25, 26^ and they have a heightened threshold for perceiving phosphenes induced by electrical stimulation of the eye, consistent with interference from spontaneous retinal activity^27^. However, without tools to suppress hyperactivity, it is impossible to differentiate the direct effects of photoreceptor signal loss from the indirect effects of increased RGC background firing (i.e.: “noise”).

Here we have used a pharmacological approach to block RA signaling, allowing us to discriminate the effects of decreased signal from increased noise on vision in the rd10 mouse. By investigating perception with visual behavioral experiments and higher-order neural processing with functional imaging of cortex, we show that RA-induced retinal hyperactivity is a major contributor to vision impairment. Moreover, by inhibiting RA, we reveal a new therapeutic strategy for mitigating vision loss that may be applicable across a wide range of photoreceptor degenerative disorders, regardless of the underlying etiology.

## Results

### Inhibitors of RA synthesis or RA signal transduction reduce degeneration-induced gene expression in the retina of rd10 mice

We intervened at two different steps in the RA signaling pathway in an attempt to suppress or reverse degeneration-induced remodeling of RGCs in the rd10 mouse model of RP (**Fig. 1A**). RA is a retinoid, derived from dietary retinol (vitamin A). Retinol is converted into retinaldehyde by retinol dehydrogenase (RDH), which is expressed in retinal pigment epithelium (RPE) cells and Müller glial cells^28–30^. 11-cis retinaldehyde, the opsin chromophore for rod and cone phototransduction, can be regenerated after photoisomerization by enzymes of the visual cycle, which are also expressed in RPE and Müller glial cells. However, retinaldehyde can also be converted into RA by the enzyme retinaldehyde dehydrogenase (RALDH), which is expressed in choroid, RPE, as well as retina^31, 32^. RA crosses cell membranes and binds to RAR, a nuclear protein that heterodimerizes with the retinoid orphan receptor (RXR). The complex, in conjunction with other co-activator proteins, then binds to specific DNA sequences to enhance transcription of genes^33–35^. RAR and RXR are expressed in many cells, but since physiological changes intrinsic to RGCs can account for much of degeneration-induced hyperactivity^17^ we focused specifically on RGCs.

**Figure 1:**
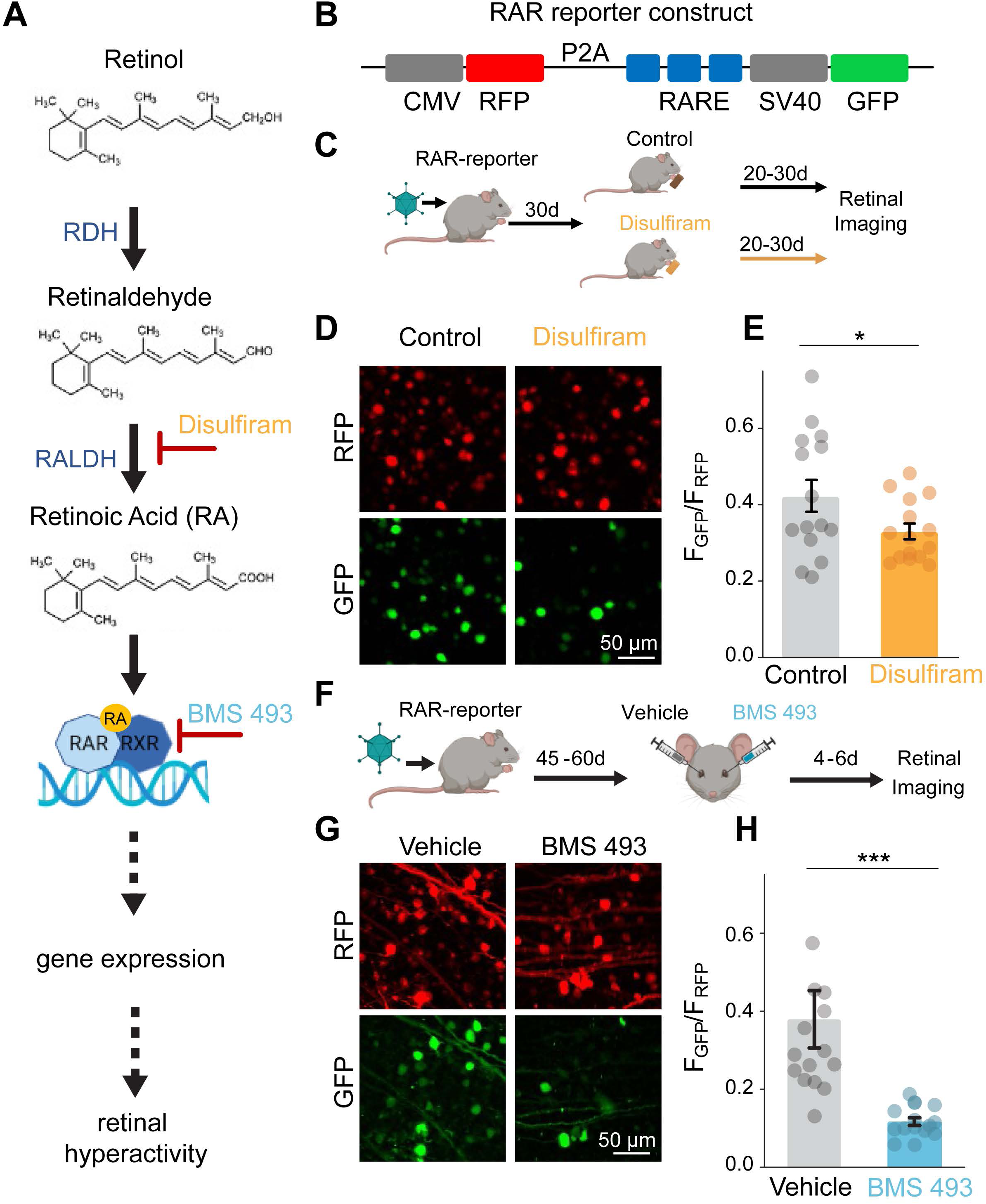
Disulfiram and BMS 493 reduce signaling through the RA pathway in the degenerated retina. **A**) RA signaling pathway, including sites of intervention by disulfiram and BMS 493. **B**) Design of the RAR-reporter. CMV: cytomegalovirus promoter (constitutive expression); RFP: red fluorescent protein; P2A: 2A self-cleaving peptide; RARE: RA-response element (RA-dependent enhancer); SV40: simian virus 40 promoter (weak expression); GFP: green fluorescent protein. **C**) Design of the disulfiram experiment. AAV2 encoding the RAR-reporter was injected into both eyes of rd10 mice at P30. Starting at P60, mice received food with disulfiram or no drug for 20-30 days. Retinas were obtained at P80-90 for imaging. **D**) Flattened Z-stacks of the ganglion cell layer showing cells expressing RFP (red) and GFP (green), obtained with a spinning-disk confocal microscope. **E**) Quantification of RA-induced expression. The value of GFP/RFP was measured for each individual labeled cell. GFP/RFP values were averaged across the many cells in each retinal sample (circles). Bar graph show the mean values for each experimental group ± SEM (n=15 retinal samples / condition in a total of 6 mice; *p<0.05, 1-tailed t-test, **Table S1**). **F**) Design of the BMS 493 experiment. AAV2 encoding the RAR- reporter was injected into both eyes at P30. At 45-60 days later, one eye was injected with 1 μl vehicle (PBS x1) and the contralateral eye with 1 μl BMS 493 (5 μM). Retinas were obtained 4-6 days post-injection and used for imaging. **G**) Representative images for **F**. **H**) Quantification of RA-induced gene expression similar to **E**. n=15 retinal samples in a total of 8 eyes from 4 mice; ***p<0.00001, Mann-Whitney test, see **Table S1**.

To assess pharmacological inhibition of degeneration-induced gene expression, we used a reporter gene construct delivered to RGCs with an adeno-associated virus (AAV) vector (**Fig. 1B**). The construct encodes red fluorescent protein (RFP), which is expressed constitutively in all virally transduced cells; and green fluorescent protein (GFP), which is expressed only upon RAR activation by RA^17, 23^. Since GFP expression reflects the level of RAR activation and RFP expression reflects the level of viral transduction that can vary between retinas, the GFP/RFP ratio can be used as a metric of degeneration-induced RA signaling^17^.

Our first target for intervention was RALDH. RALDH is a member of a large family of enzymes known as aldehyde dehydrogenases (ALDHs). Disulfiram (Antabuse®) is an FDA-approved irreversible inhibitor of ALDHs^36^, including all members of the RALDH subfamily^37^. Disulfiram is usually prescribed for chronic alcoholism and is taken orally. Ingested ethanol is converted into acetaldehyde, which is broken down by ALDHs. By preventing its breakdown, disulfiram allows buildup of acetaldehyde, discouraging alcohol consumption by causing severe hangover symptoms, known collectively as the disulfiram ethanol reaction^38^. In the absence of alcohol use, treatment with disulfiram causes infrequent adverse reactions that resolve completely after stopping treatment^39^.

To assess whether disulfiram can inhibit degeneration-dependent activation of the RA pathway, we injected the eyes of rd10 mice with the RAR-reporter virus early in degeneration (**Fig. 1C**). We continuously provided them with *ad libitum* regular food or food containing disulfiram (2 mg/Kg) for 20-30 days and imaged their retinas later in degeneration (**Fig. 1D**). RGCs in disulfiram-treated mice showed a GFP/RFP of 0.33±0.02, significantly lower than control mice with a ratio of 0.42±0.04 (p=0.028, 1-tailed T-test) (**Fig. 1E, Table S1**). These findings establish that orally administered disulfiram can be absorbed in the gastrointestinal tract, cross the blood-retina barrier, and reach the retina at a concentration sufficient to suppress RA-induced gene expression.

Our second target for intervention was RAR. We used BMS 493, a high-affinity inverse agonist of all RAR isoforms^40^. We showed that BMS 493 increases the response of RGCs to photostimulation of residual rods and cones in rd10 mice^17^, supporting our conclusion that RA, signaling through RAR, mediates degeneration-induced physiological remodeling.

To test whether BMS 493 can inhibit degeneration-dependent activation of the RA pathway, we injected rd10 mice with the RAR-reporter virus early in degeneration (**Fig. 1F**). In each mouse, we delivered BMS 493 in one eye and vehicle in the contralateral eye, by intravitreal injections. Finally, we imaged the retinas isolated from these mice (**Fig. 1G**). RGCs in BMS 493-injected eyes had a mean GFP/RFP value of 0.12±0.01, significantly lower than RGCs in vehicle-injected eyes with a ratio of 0.38±0.07 (p<0.00001, Mann-Whitney test) (**Fig. 1H, Table S1**). These findings establish that delivery of BMS 493 through intravitreal injection suppresses RA-induced gene expression even more effectively than orally administered disulfiram.

### Disulfiram and BMS 493 reduce RA-induced RGC hyperactivity without affecting residual photoreceptor function

We next asked if inhibiting the RA pathway can suppress RGC hyperactivity in the rd10 retina. We used mice mid-way through the photoreceptor degeneration process (P40-45) when RGCs are already hyperactive^17, 41^. Oral disulfiram must cross several barriers to inhibit RA synthesis and subsequent downstream effects in the retina, so we assessed its effects on hyperactivity after 40 days of continuous treatment (**Fig. 2A**), sufficient time to significantly inhibit RA-induced gene expression (**Fig. 1E**). Control littermates were given food without disulfiram for the same time period.

**Figure 2:**
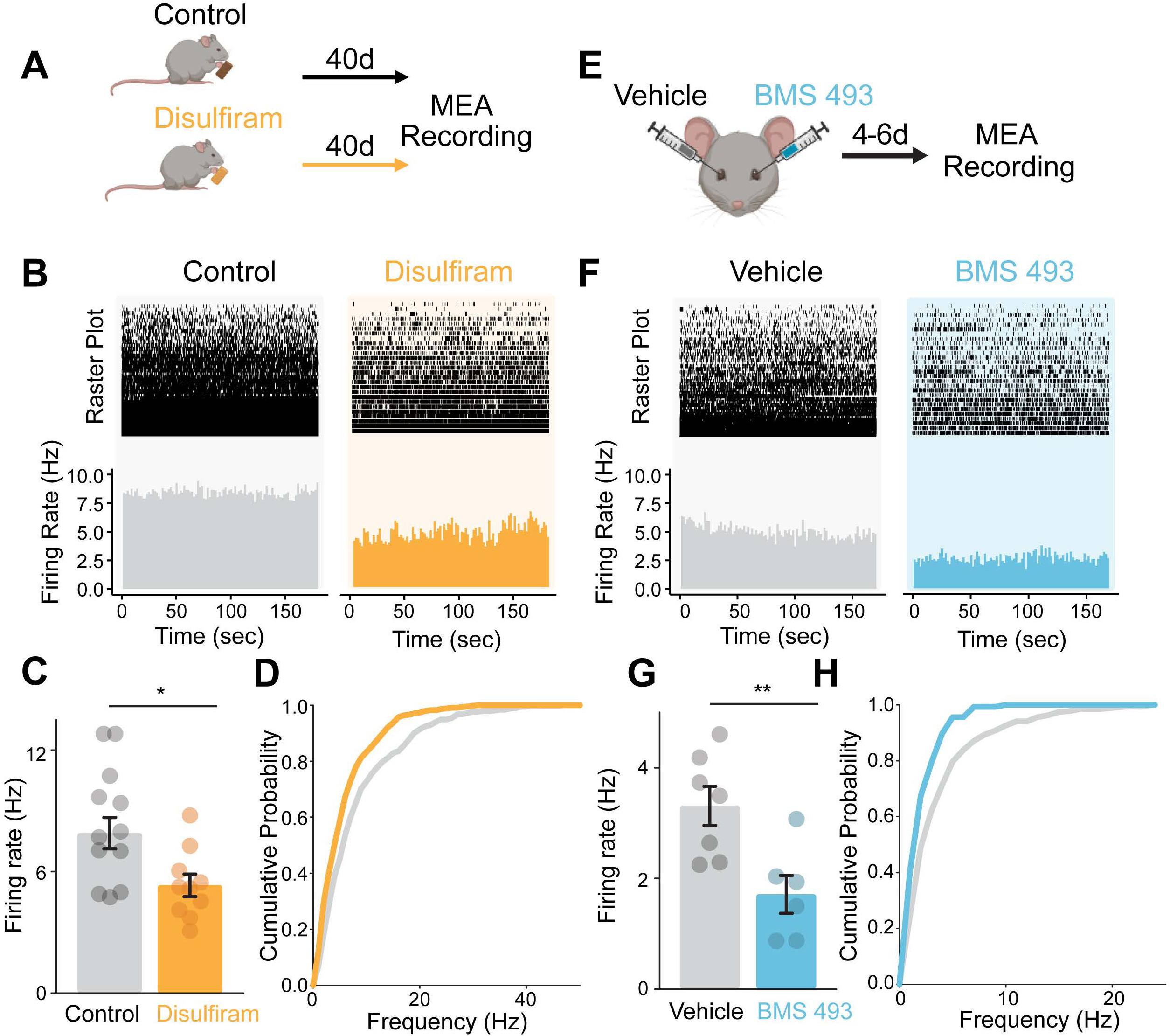
Disulfiram and BMS 493 reduce hyperactivity in the degenerating retina. **A)** Design of the disulfiram experiment. Early degeneration rd10 mice (P40-45) were given food with or without 2 mg/Kg disulfiram for 40 days. At P80-P85, retinas were obtained and prepared for MEA recording of spontaneous RGC activity in darkness. **B)** MEA recording from retinal samples (∼15 mm^2^ each) from control and disulfiram mice. Raster plots show spontaneous activity of individual units (∼60 each). The bottom panels show mean firing rates across all the units. **C)** Mean firing frequency for all RGCs in each individual retinal sample (circles) and across the entire set of samples from each group (bar graphs). Values are shown as mean ± SEM, *p<0.05, Mann-Whitney test (**Table S2**). **D)** Cumulative probability of firing frequencies in samples from control and disulfiram retinas (control: n=568 units, 11 retinal samples from 4 mice; disulfiram: n=443 units, 10 retinal samples from 3 mice). Disulfiram (orange curve) resulted in a leftward shift in the cumulative probability curve, indicating reduced firing across all units (p<0.001, Kolmogorov–Smirnov test, **Table S2**). **E)** Design of the BMS 493 experiment. Early degeneration rd10 mice (P40-P45) were injected intravitreally with 1 μl vehicle (PBS x1) in one eye, and 1 μl BMS 493 (5 μM) in the other eye. At 4-6 days post-injection, retinas were obtained and prepared for MEA recording of spontaneous RGC activity in darkness. **F)** Retinal MEA recordings from vehicle and contralateral BMS-493 injected eyes from one individual mouse. **G)** Mean firing frequency similar to **C**, **p<0.01, Mann-Whitney test (**Table S2**). **H)** Analysis of spontaneous activity, similar to **D**. Cumulative probability plot of firing frequencies in the active units of vehicle and BMS 493-injected eyes (vehicle: n=271 units from 7 retinal samples; BMS 493: n=135 units in 6 retinal samples; a total of 6 eyes from 3 mice). The left shift in the BMS 493 curve (light blue) is statistically significant (p=0.001, Kolmogorov–Smirnov test, **Table S2**).

After treatment, we isolated the retina and prepared flat-mounted samples for multielectrode array (MEA) recordings of spontaneous RGC activity. Recordings showed less spontaneous firing in darkness in samples from disulfiram-treated mice than from control mice (**Fig. 2B**). Overall, mean firing frequency across all units was 5.34±0.54 Hz with disulfiram, significantly less than 7.90±0.78 Hz for control (p=0.016, Mann-Whitney test) (**Fig. 2C, Table S2).** Cumulative probability analysis shows that disulfiram scales down firing in highly-active units (**Fig. 2D**, p<0.001, Kolmogorov– Smirnov test). In disulfiram-treated retinas, we detected a mean value of 44.3 spontaneously active units per retinal sample, as compared to 51.6 active units in control retinas, a reduction of 14.2% (**Table S2**).

Next, we assessed BMS 493 (**Fig. 2 E-H**). One eye of each mouse was injected with BMS 493 and the contralateral eye with vehicle, allowing intra-animal comparisons (similar to **Fig. 1F**). We again used mid-degeneration rd10 mice (P40-45), but because the drug was injected directly into the eye, and administered only once, inhibition of RA signaling begins much more quickly and is more transient. Therefore, recordings were obtained much earlier in the life of the mice, at 4-6 days after injection (P45-50) (**Fig. 2E**). This is long enough to significantly inhibit RA-induced gene expression (**Fig. 1H**), but at a younger total age than in the disulfiram experiments, accounting for the lower baseline firing rate (compare **Fig. 2, B** vs **F**).

Retinas obtained from eyes injected with BMS 493 showed less spontaneous activity than retinas from contralateral eyes injected with vehicle (**Fig. 2F**). The mean firing rate of BMS 493-treated retinas was 1.71±0.34 Hz, significantly lower than in vehicle-injected retinas of 3.31±0.35 Hz (**Fig. 2G**, p<0.006, Mann-Whitney test). Cumulative probability analysis of all units shows that like disulfiram, BMS 493 decreased the probability of encountering high firing frequency RGCs (**Fig. 2H**, p<0.001, Kolmogorov– Smirnov test). In BMS 493-treated retinas, the mean number of spontaneously active units per retinal sample was 22.5, while the same value was 38.7 in the vehicle-injected counterpart eye of the same mouse, a decrease of 41.9% (**Table S2**). This result also suggests that the intravitreal injections of vehicle (PBS x1) do not cause further degeneration that can lead to hyperactivity.

RAR expression is abundant in the outer retina, particularly in cone photoreceptors^42^. Hence it seemed possible that disulfiram and BMS 493 might affect events in the outer retina, independent of their effect on RGCs. To address this possibility, we recorded the photopic electroretinogram (ERG) in rd10 mice^43^. We focused on the b-wave, which reflects light-modulated synaptic transmission primarily from cones to bipolar cells, since the rods are largely degenerated at this age (P60-80). We found that neither disulfiram (**Fig. S1A,B**) nor BMS 493 (**Fig. S1C-E**) caused a significant change in the amplitude of the b-wave across a range of photopic intensities (see also **Table S3**).

These results suggest that inhibiting RA does not affect events in the outer retina, leaving events in the inner retina as the primary mechanism of RA-induced hyperactivity.

### Disulfiram and BMS 493 enhance behavioral detection of images in rd10 mice

Genetically interfering with RA signaling augments light-avoidance behavior in rd10 mice^17^, but whether inhibiting RA signaling can improve image-forming vision has remained unknown. To investigate this, we used a visual detection test involving operant conditioning, inspired by a previous paradigm for measuring visual threshold^44^. We used a behavioral cage outfitted with a unidirectional running wheel, orienting the mouse towards a computer display (**Fig. 3A**). The cage also has a nose poke sensor, which when engaged triggers delivery of a water droplet reward (10% sucrose).

**Figure 3:**
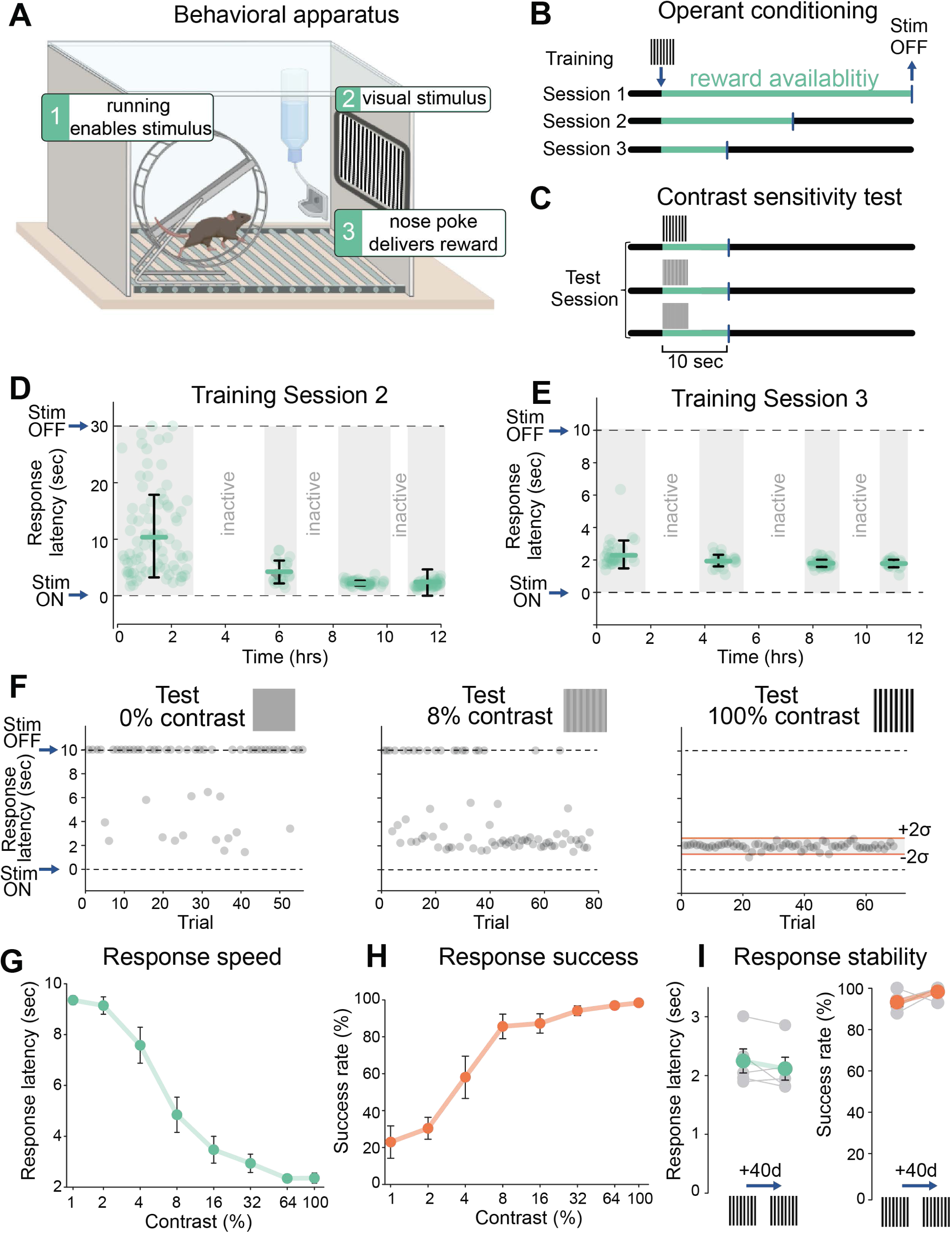
Measuring behavioral image detection in mice. **A)** Apparatus for operant-conditioned image detection. Wheel running gates the display of a conditioned stimulus image associated with the availability of reward (a 10% sucrose water droplet). Reward is obtained by engaging the nosepoke within the allotted reward interval. **B)** Training to 100% contrast images occurs over 3 sessions (each 12 hours) with decreasing reward intervals, shaping response latency. (**Table S4**). **C)** Contrast test was performed under the same conditions as session 3, but the visual stimulus presented was randomized between discrete contrasts from 0-100%. See **Table S4**, **Fig. S2**. **D-F)** Responses of an individual wild-type mouse during training session 2 (**D**) and 3 (**E**), and to 3 different contrasts (0%, 8% and 100%) during contrast sensitivity test (**F**). Trials of equal contrasts were grouped for quantifying latency, even though they were randomized during the test. Red lines in responses to 100% contrast show the upper and lower limits of the value range for a single trial to be considered successful (mean±2SD). **G)** Grouped data showing full contrast-sensitivity curve. The response speed (latency in seconds) plotted against the contrast of each visual stimulus (n=5 WT mice, P60). **H)** Full contrast sensitivity curve showing success rate as a function of contrast in the same group of mice as in **G**. **I)** Comparison of response speed and success rate in the same group of wild-type mice at P60 vs P100, with no training session carried out between the two time points. Individual data for each mouse shown in grey, mean ± SEM values shown in green (latency) and red (success rate), respectively. p>0.05, Wilcoxon rank sum test.

Running on the wheel digitally gates the onset of a full-contrast grating displayed on the screen, while simultaneously unlocking the nose poke. Mice are water- and food-deprived before each session, providing motivation for discovering the association between the visual stimulus and availability of reward. The period of reward availability was shortened during 3 successive training sessions to increase response stringency (**Fig. 3B, Table S4**). Only those mice achieving a criterion level of success during training (>80% of trials with response latency <4 sec) with 100% contrast were carried forward for subsequent testing of contrast sensitivity. This involved randomly varying the image contrast over 9 distinct values over the course of a testing session (**Fig. 3C**). All images were equi-luminant irrespective of contrast, and monitors across different cages were calibrated for uniform brightness (see **Materials and Methods**).

**Fig. 3D-F** exemplifies the behavioral responses of an individual wild-type mouse during training and testing. During training, the mouse learns to associate the full-contrast image with the availability of reward (**Fig. 3D**). At the beginning of training session 2, response latencies were prolonged and inconsistent, but by the end, responses occurred within 2-3 seconds of stimulus presentation. The response latency remained at 2-3 seconds during training session 3 when the reward availability period was reduced (**Fig. 3E**), further reinforcing the association.

We tested contrast sensitivity on a mouse that had achieved the criterion level of success during training. We presented stimulus images at different contrasts, randomized from trial to trial (**Fig. S2**). Low contrast images elicited responses with longer and more variable latencies than high contrast images (**Fig. 3F**). The mean latency of 5 WT mice subjected to the contrast sensitivity test is gradually reduced from ∼8 seconds at 1% contrast to ∼2 seconds at 100% contrast (**Fig. 3G**). The mean latency at 100% contrast was used to set the range of responses considered as successful ones for all other contrasts and subsequent tests (**Fig. 3F, ‘Test 100% contrast’**). A nose poke was considered successful if it happened at a time range of -2 standard deviations (SD) to +2SD of the mean (a mouse with a response SD ≥ 40% of the mean was excluded from analysis, see **Materials and Methods** and **Table S5,S6**).

Success rate was then used to generate a full contrast-sensitivity curve (**Fig. 3H**). To test the possibility that mice might fail the test due to behavioral extinction, we tested the same 5 mice 40 days apart with no ‘refresher’ training session in between (**Fig. 3I**). We found that their response latency and success rate did not differ in the second test from the first, demonstrating that, once they learn, mice are able to retain this visually guided operant conditioning for long periods of time.

Next, we carried the same procedure to quantify the effects of disulfiram on contrast-sensitivity. Mice younger than P35 run on the wheel infrequently and are poor learners of this task. Therefore, we started training at P35-40, when some photoreceptor degeneration had already begun (**Fig. 4A**), but early enough to ensure consistent short-latency responses to the 100% contrast image (**Fig. 4B,C, left**). All mice were first trained and tested in the absence of any pharmacological manipulation and were randomly assigned to treatment or control groups ahead of the second test.

**Figure 4:**
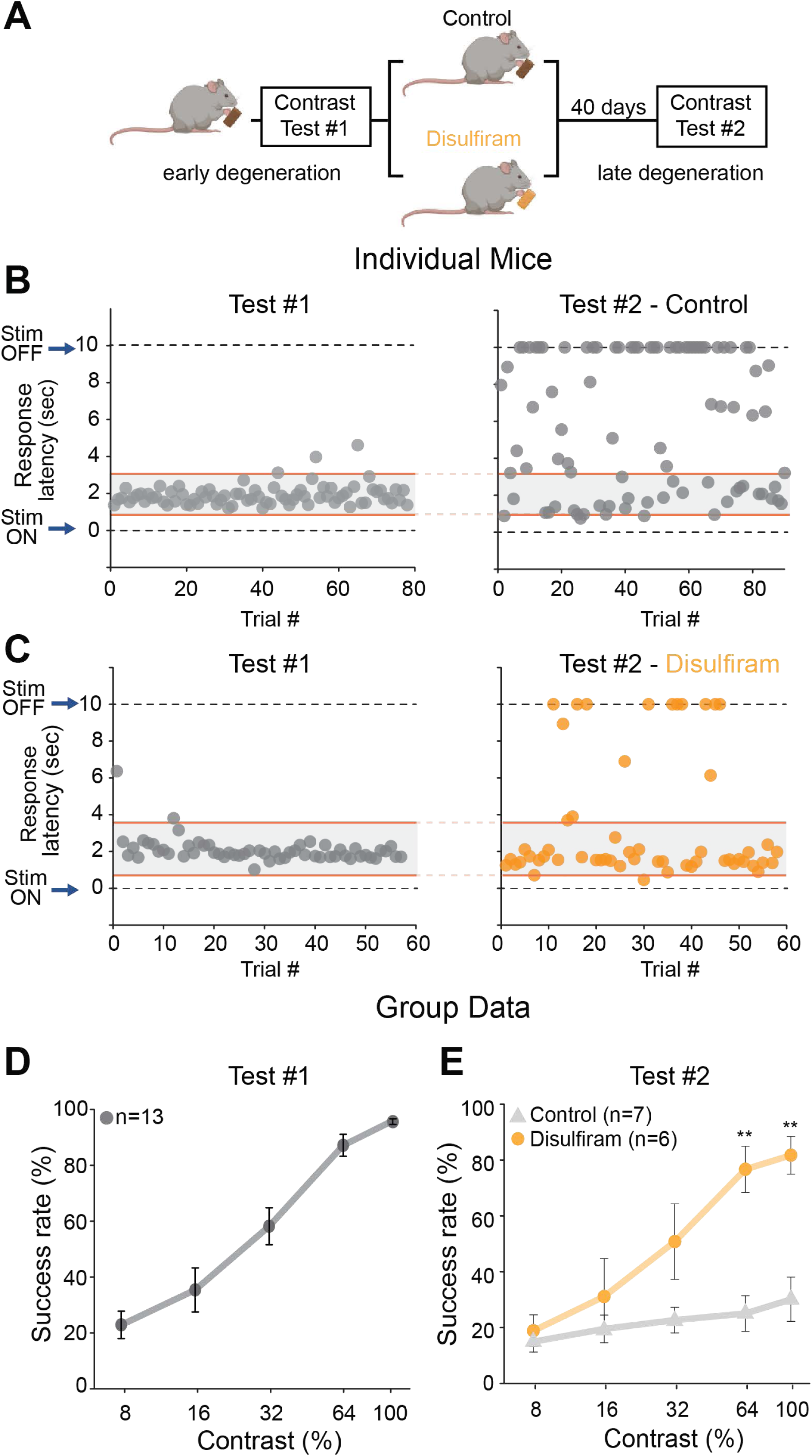
Disulfiram, a RALDH inhibitor, improves visual contrast sensitivity in rd10 mice. **A**) Experimental design. Rd10 mice were trained and tested at P35-40 (test #1), randomly separated into control or disulfiram groups, and tested again 40 days later at P75-80 (test #2). Training session 3 (Fig. 3B) was used as “refresher” training every ∼10 days. **B,C**) Responses of individual mice in the control (**B**) and disulfiram (**C**) group. The response speed (latency) to the visual stimulus at 100% contrast is shown for each individual trial, at early degeneration and before intervention (test #1, left) and at late degeneration and after intervention (test #2, right). **D**) Results of 13 mice before on test #1. Success rate is plotted as a function of contrast, values are shown as mean ± SEM (**Table S5**). **E**) Contrast sensitivity curve obtained later in degeneration and following 40 days of treatment with disulfiram or no treatment in control mice. **p<0.01, Mann-Whitney test (**Table S5**).

**Fig. 4B** shows the behavior of a trained rd10 mouse early (test #1) and late (test #2) in degeneration. Without drug treatment, short-latency responses disappeared nearly completely due to vision loss. However, in a mouse receiving disulfiram, short-latency responses were retained (**Fig. 4C**). Group data from early degeneration rd10 mice shows the already impaired contrast sensitivity caused by partial death of photoreceptors (compare **Fig. 4D** to **Fig. 3H**). After test #1 was complete, mice were randomly assigned to the control or disulfiram group, but retrospective analysis of their performance in test #1 showed similar contrast sensitivity (p>0.05, Mann-Whitney test, **Table S5**). However, when tested for the second time 40 days later, mice fed with disulfiram-containing food (2 mg/Kg) performed significantly better than control when reacting to high-contrast images (64% and 100% contrast, p<0.01, Mann-Whitney test, **Table S5**). As expected, experiments on wild-type mice showed no significant effect of disulfiram on behavioral contrast sensitivity (**Fig. S3A**), consistent with disulfiram acting specifically by inhibiting RA synthesis.

A similar series of tests was conducted to assess the efficacy of BMS 493 in boosting contrast sensitivity in degenerative rd10 mice (**Fig. 5A**). Similar to experiments with disulfiram (**Fig. 4**), rd10 mice were trained and tested, and then randomly assigned for binocular intravitreal injections of BMS 493 or vehicle. An individual rd10 mouse receiving intravitreal BMS 493 showed more short latency responses than an age-matched rd10 mouse injected with vehicle (**Fig. 5B,C**). Contrast-sensitivity curves show that BMS 493 treatment preserved short-latency responses to 100% contrast stimuli (**Fig. 5D,E**) while almost none of these responses were seen in vehicle-treated mice (p<0.05, Mann-Whitney test, **Table S6**). Control experiments on wild-type mice showed no augmentation of vision after BMS 493 treatment (**Fig. S3B**). This is consistent with our previous studies showing little or no ongoing RA-induced gene expression in healthy retinas of wild-type mice^17^, supporting the conclusion that BMS 493 improves vision specifically by blocking RAR. Taken together, these findings indicate that behavioral vision loss can be alleviated by inhibiting the RA pathway, consistent with RA-induced remodeling making a major contribution to corrupting vision in retinal degenerative disorders.

**Figure 5:**
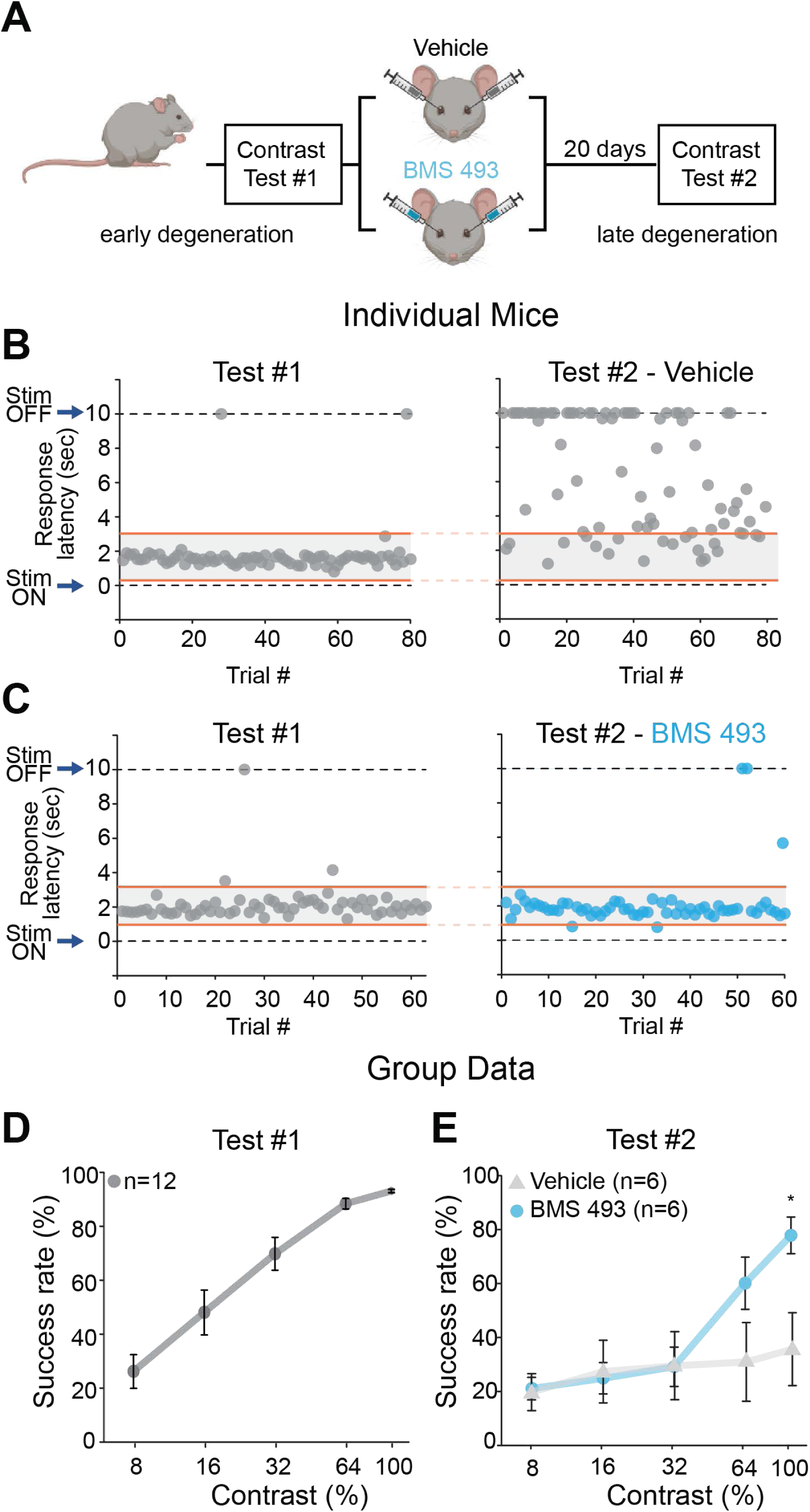
BMS 493, an RAR inhibitor, improves visual contrast sensitivity in rd10 mice. **A)** Experimental design. Rd10 mice were reared in the dark until P55. At P55-60, they were trained and tested for contrast sensitivity and placed under 14:10 L/D conditions to induce degeneration. They received binocular intravitreal injections containing 1 μl of BMS 493 (5 μM) or vehicle (PBS x1), and tested for a second time 4-6 days later, at P80-85. **B,C**) Responses of individual mice in the vehicle (**B**) and BMS 493 (**C**) group. The response speed (latency) to the visual stimulus at 100% contrast is shown for each individual trial, at early degeneration and before intervention (test #1, left), and at late degeneration and after intervention (test #2, right). **D**) Results of 6 vehicle mice (grey) and 6 BMS 493 mice (blue) before administration of drug during test #1. Success rate is plotted as a function of contrast, values are shown as mean ± SEM, p>0.05 for all contrasts tested (**Table S6**). **E**) Contrast sensitivity curve obtained later in degeneration and following treatment with vehicle or BMS 493. *p<0.05, Mann-Whitney test (**Table S6**).

### Inhibiting the RA pathway sharpens the tuning of cortical neurons to complex visual stimuli

Behavioral evaluation of image-forming vision in mice requires days of training and testing, but physiological evaluation of neuronal responses can be characterized rapidly, without behavioral training, through functional Ca^2+^ imaging. Calcium imaging has the additional advantage of providing responses from many neurons simultaneously. Responses to a wide range of visual stimuli, from bars of specific orientation to movies with complex naturalistic scenes, can be elicited in a single session. Cortical imaging can provide a more thorough description of how photoreceptor degeneration degrades visual function and how the RA pathway inhibitors disulfiram and BMS 493 might alleviate vision impairment.

To test the effect of disulfiram, rd10 mice were given a diet with or without the drug for 28 days, beginning early in the process of degeneration and ending late, after visual function was severely impaired (**Fig. 6A**). Midway through the treatment period we performed a surgical procedure (a hemicraniectomy) under anesthesia to facilitate intracortical injection of an AAV encoding the Ca^2+^ indicator GCaMP6s, which included the Ca^2+^/calmodulin-dependent protein kinase II (CaMKII) promoter to target expression to excitatory neurons (see **Materials and Methods**). Following injection, we inserted a glass cranial window to provide optical access for subsequent imaging of GCaMP6s expressing neurons at the end of the treatment period. Imaging data were collected blind to treatment conditions.

**Figure 6:**
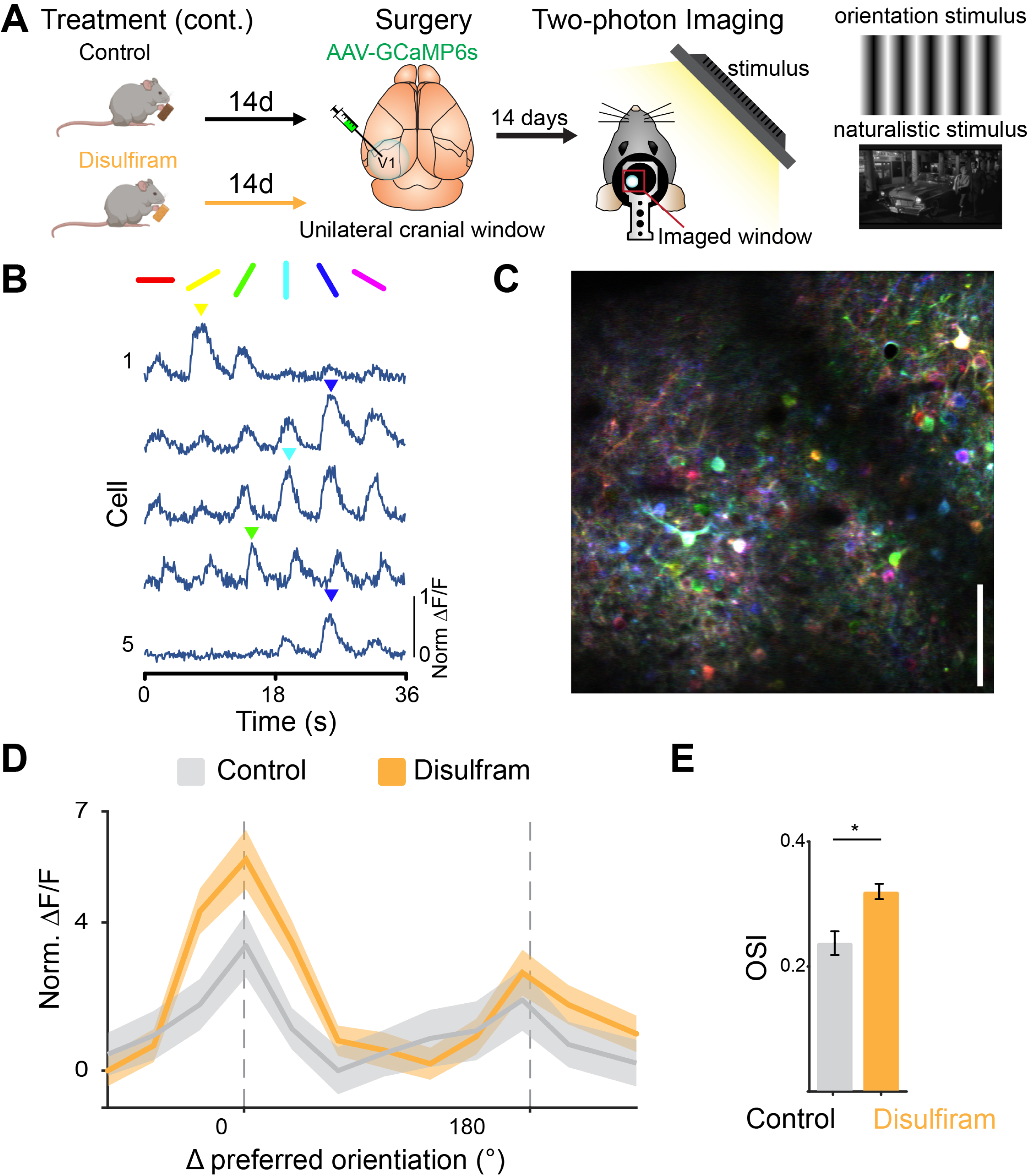
Disulfiram treatment sharpens orientation coding of visual cortical neurons. **A)** Experimental timeline for disulfiram experiments. P21 rd10 mice were placed on either a control or disulfiram diet for 14 days prior to virus injection and cranial window implantation at P35. After 14 additional days (P49), mice were imaged under a two-photon microscope during presentation of visual stimuli, either oriented gratings or naturalistic scenes, to the contralateral eye. **B)** Example traces from individual neurons in response to the drifting grating stimulus, showing responses to specific orientations. **C)** Example imaging field from the two-photon microscope, with neurons pseudo colored by preferred orientation (color map in right panel). Scale bar = 100 microns. Note that neurons that respond to all gratings in a non-selective manner appear white.**D)** Aligned and averaged cross-validated tuning curves across neurons in each condition. Preferred orientation is determined using odd trials and averaged responses are shown for even trials. Disulfiram treatment results in increased response amplitude at the preferred orientation as compared to control. **E)** Disulfiram treated neurons have significantly greater orientation selectivity indices (OSI) than control neurons. Values shown are mean ± SEM. *p<0.05, 2-tailed t-test.

Many excitatory neurons in the primary visual cortex respond selectively to stimuli of a particular orientation^45^. To determine whether disulfiram treatment influences orientation tuning, we measured responses to drifting periodic gratings presented to the eye contralateral to the cranial window (**Fig. 6B,C**). We found that a lower percentage of neurons were orientation-tuned in rd10 mice (16.06%) than typically observed in wild-type mice using a similar approach (∼30% in^46^), but the fraction tuned was significantly increased with disulfiram treatment (22.69%). In addition, disulfiram treatment caused significantly sharper orientation tuning as compared to control (t(1127) = 2.01, p = 0.04; linear mixed-effects model: fixed effect for treatment, random effect for mouse; **Fig. 6D,E**). This was quantified with a cross-validated orientation selectivity index (OSI) measure that compares the magnitude of response to preferred vs. non-preferred orientations, where the preferred orientation is determined using a separate subset of trials to eliminate spurious orientation tuning due to bursts of activity (see **Materials and Methods**).

We next asked whether inhibiting the RA pathway can improve the detection of responses to complex naturalistic stimuli. To investigate this, we presented a 30 second natural scene (http://observatory.brain-map.org/visualcoding/stimulus/natural_movies) that contains a rich diversity of visual information. Others have found that individual neurons in the mouse visual cortex respond preferentially at different time points in the movie, reflecting their tuning to different combinations of higher order features^47^. By presenting the same clip to the mouse many times, we could measure the response reliability across trials, while remaining unbiased to any specific visual feature (**Fig. 7A**). A highly reliable response suggests that a neuron is coding information about that particular frame with high fidelity.

**Figure 7:**
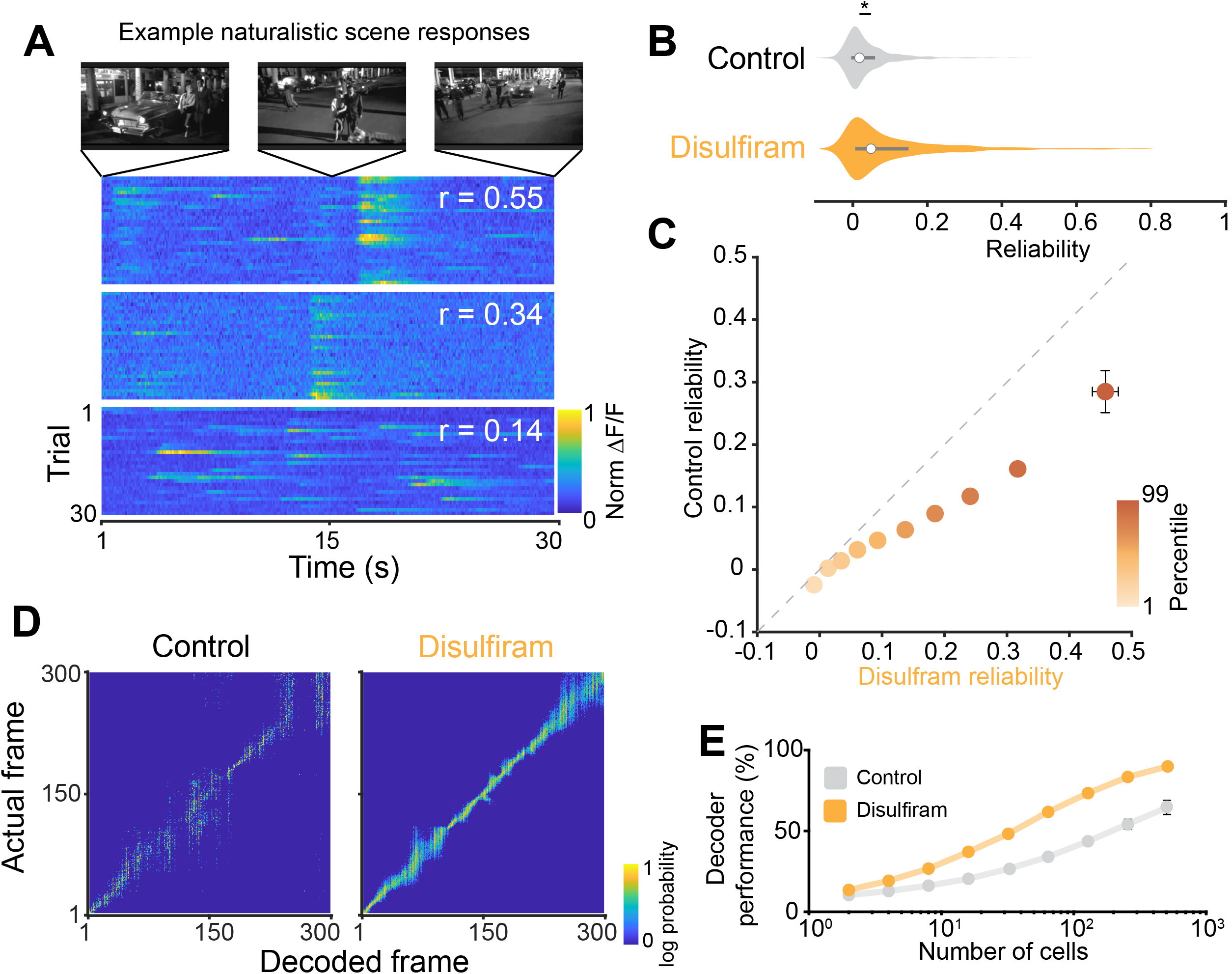
Disulfiram treatment improves coding of naturalistic scenes in V1. **A)** Top: example frames from the movie clip presented to rd10 mice. Bottom: Example responses from three different neurons to repeated presentations of the movie. Each row is the response to a single presentation. Different neurons show varying amounts of response reliability to repeated presentations. **B)** Comparison of the response reliability of neurons from disulfiram-treated versus control-treated mice (experimental design similar to Fig. 6A). Neurons in disulfiram mice show significantly greater reliability compared to neurons from control mice. **C)** Comparison of reliability segmented by deciles. Reliability is similar in lower deciles, but becomes significantly greater for disulfiram treated mice in higher deciles. **D)** Performance of a population level decoder across control (left) vs disulfiram treated (right) mice. **E)** Across all population sizes used for decoding, disulfiram treatment shows significantly improved performance (predicted frame within 10 frames of actual frame) compared to control. **B,C,E**) Values shown are mean ± SEM. *p<0.05, 2-tailed t-test.

We found high variability in the response properties of cortical neurons in rd10 mice, with many exhibiting low reliability. However, the distribution of reliability values was shifted significantly by disulfiram treatment, with an increase in the proportion of neurons showing high reliability (t(2068) = 2.14, p = 0.03; linear mixed-effects model; fixed effect for treatment, random effect for mouse; **Fig. 7B).** On average, reliability across the population was increased >1.7-fold, from 0.1022 to 0.1751 by disulfiram, consistent with higher-fidelity detection of high-order visual features. Because of the high variability in response properties, we questioned whether disulfiram affected all cells equally (an additive shift) or unequally (a multiplicative shift). We separated the data from each recording by deciles and compared the values of reliability within each decile across control and disulfiram groups (**Fig. 7C**). Overall, disulfiram treatment improves coding of natural scenes across decile groups (treatment main effect: t(131) = 4.37, p = 2.47 x 10^-5; linear mixed-effects model, fixed effect for treatment and decile, random effect for mouse). Additionally, there was a significant interaction between treatment condition and decile group, suggesting that the effects of disulfiram treatment differentially affect neurons, depending on their response fidelity (treatment and decile group interaction: t(131) = 2.99, p = 0.003; linear mixed-effects model, fixed effect for treatment and decile, random effect for mouse).

We next reasoned that sharpened feature selectivity would improve the population-level representation of the naturalistic movie stimulus. In order to test this, we used an ideal observer analysis to decode the current frame of the movie based on the neuronal population responses (**Fig. 7D**; see **Materials and Methods**). We carried out this analysis over many iterations, using different sized pools of randomly selected neurons from both disulfiram and control mice. To quantify the performance of the decoder, we calculated the percentage of iterations in which the decoded frame was within 10 frames (1 sec) of the actual frame (**Fig. 7E**). We found that decoding in disulfiram-treated mice consistently outperformed that of control mice across neuronal pool sizes (treatment group effect: t(79) = 3.95, p = 1.68 x 10^-4; linear mixed-effects model; fixed effect for treatment and pool size). These experiments indicate that disulfiram treatment leads to more robust encoding of complex features found in naturalistic scenes, consistent with improved visual perception.

We next repeated these analyses to ask whether inhibiting RAR with BMS 493 could also improve visual processing of complex stimuli (**Fig. 8A**). For these experiments, each mouse was anesthetized and BMS 493 was injected intravitreally into one eye while the contralateral eye received vehicle. At 4-6 days after injection, we compared responses of cortical neurons to visual stimuli displayed individually to each eye. We recorded Ca^2+^ responses from the monocular zone of the primary visual cortex, which receives information selectively from the contralateral eye. This procedure allowed a within-mouse comparison of cortical response properties, eliminating possible concerns about variability in the degree of retinal degeneration between individual mice.

**Figure 8:**
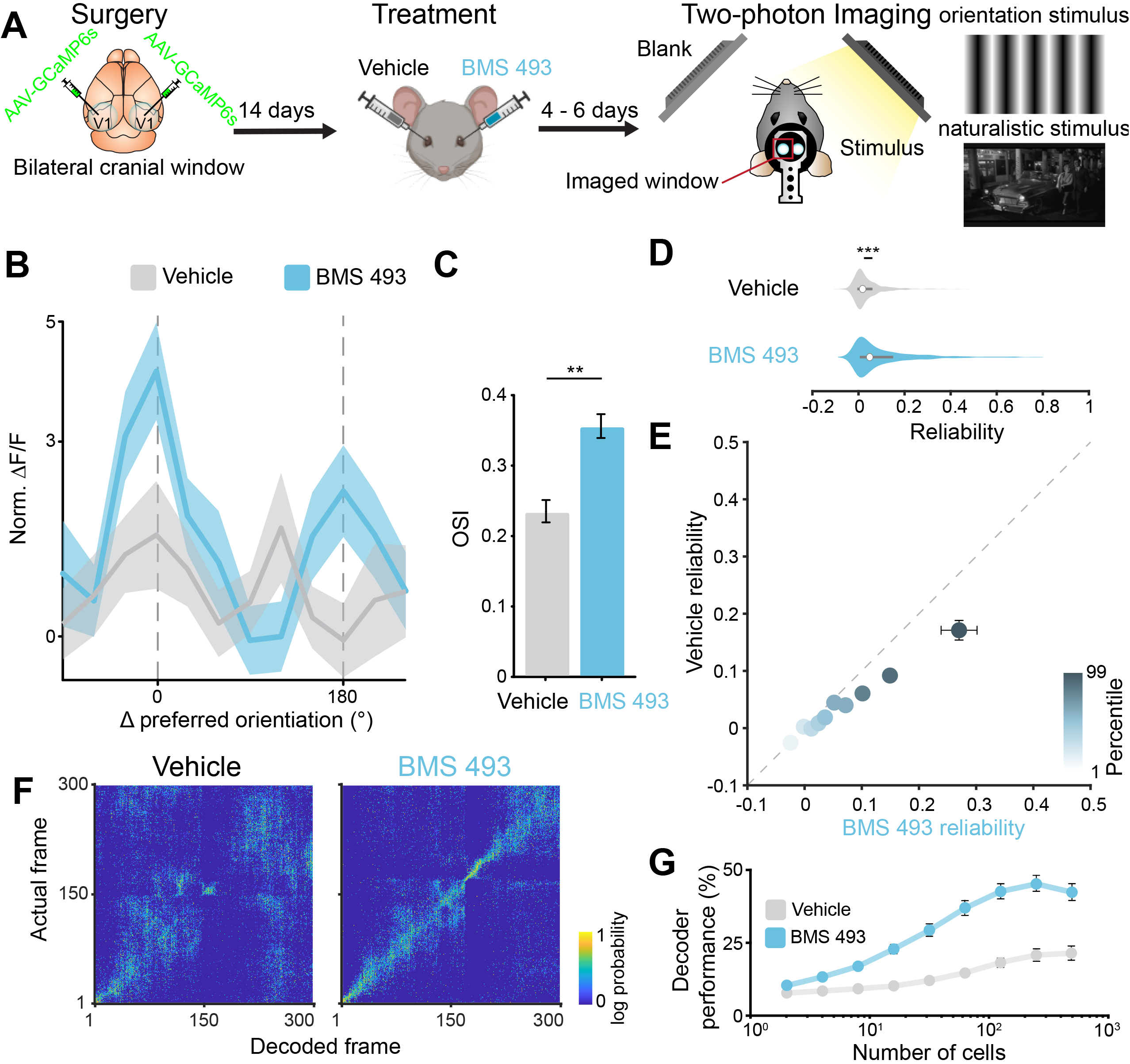
Intravitreal BMS 493 injection improves visual processing in contralateral V1. **A)** Experimental timeline for BMS 493 experiments. P28 rd10 mice were given a bilateral cranial window and allowed to recover for 14 days prior to treatment. At P43-45, vehicle or BMS 493 were injected into separate eyes. Mice were imaged 4 -6 days post-treatment at P49. This allowed comparison of drug effects on retinal-driven cortical responses within the same mouse. **B)** Aligned and averaged cross-validated tuning curves across neurons in each treatment condition. Preferred orientation is determined using odd trials and averaged responses are shown for even trials. Injection of BMS 493 results in increased response amplitude at the preferred orientation compared to vehicle. **C)** Neurons contralateral to the BMS 493 treated eye show increased OSI as compared to neurons contralateral to the vehicle eye. **D)** Neurons contralateral to the BMS 493 treated eye show significantly higher average reliability compared to neurons contralateral to the vehicle treated eye. **E)** Comparison of reliability segmented by deciles. Reliability is similar in lower deciles, but becomes significantly greater for BMS 493 neurons at higher deciles. **F)** Performance of a population level decoder across vehicle (left) and BMS 493 treated (right) populations. **G)** Across population sizes, BMS 493 treatment exhibits improved decoding performance as compared to vehicle. C,D,E,G) Values are shown as mean ± SEM, ***p<0.001, two-tailed t-test.

Comparison across treated and untreated neuronal populations showed significantly increased OSIs in neurons receiving input from the BMS 493-treated eye than from the vehicle-treated eye (t(1078) = 3.26, p = 1.16 x 10 ^-3; linear mixed-effects model; fixed effect for treatment, random effect for mouse; **Fig. 8B,C**). These neurons also exhibited dramatically increased response reliability to the naturalistic movies (t(1961) = 10.12, p = 1.75 x 10 ^ -23; linear mixed-effects model; fixed effect for treatment, random effect for mouse; **Fig. 8D**). As with disulfiram, comparing reliabilities across deciles also suggested a multiplicative effect of BMS 493 (treatment group effect: t(206) = 4.0266, p = 7.95 x 10^-5; treatment and decile group interaction: t(206) = 2.84, p = 0.005; linear mixed-effects model, fixed effect for treatment and decile, random effect for mouse; **Fig. 8E**). Finally, ideal observer analyses showed much greater decoding accuracy in V1 neurons contralateral to the BMS 493 treated eye compared to neurons contralateral to the vehicle treated eye (treatment group effect: t(144) = 3.7669, p < 10^-5; linear mixed-effects model; fixed effect for treatment and pool size; **Fig. 8F,G**). Taken together, these experiments indicate that BMS 493 can rescue responses to complex visual stimuli in downstream cortical neurons, during retinal degeneration.

## Discussion

### Degeneration-induced retinal hyperactivity corrupts vision in rd10 mice

As photoreceptors degenerate, the retina undergoes morphological and physiological plasticity, some of which is thought to be adaptive, but much of which is maladaptive. On one hand, downstream circuit function is resilient, such that selective loss of rods leads to compensatory changes in the rod synaptic pathway to restore synaptic strength^48, 49^, while loss of cones leads to similar changes in the cone pathway^50^. On the other hand, hyperactivity triggered by RA can be maladaptive, adding background noise that obscures signals that are already attenuated by the loss of photoreceptors^17^. Our behavioral and cortical imaging results indicate that RA-induced changes in neural processing impairs not only simple light detection by the mouse, but also higher-order visual capabilities, such as reliable detection of specific visual scenes.

Visual information is ordinarily transmitted from the retina to the brain in the form of a RGC spike frequency code. Hypothetically, the code might be replicable with prosthetic devices or optogenetic tools that stimulate RGCs artificially in response to light^51^, but accurate replication of the code necessitates that the background firing rate of RGCs is reliable. If artificially stimulated firing is superimposed on hyperactive spontaneous firing, the proper encoding of input stimuli would be altered, corrupting the accurate representation of visual features. This should apply whether spikes are driven naturally by residual photoreceptors or artificially by prosthetic means. Our behavioral and cortical imaging results indicate that corrupted neural processing impairs vision at various levels of brain function and perception, from behavioral light detection^17^ and image recognition (**Figs. 4,5**) to reliable representation of complex naturalistic scenes by the visual cortex (**Figs. 6-8**). The loss of photoreceptors is the most obvious cause of vision loss, but our findings indicate that physiological remodeling in downstream neurons contributes significantly to visual decline. Moreover, inhibiting RA-induced changes can mitigate impairment of simple or complex visual functions, even in mice with late-stage photoreceptor degeneration. These findings suggest a new therapeutic avenue for improving low vision in humans.

Previous work indicated that elevated RA signaling, and its ensuing hyperactivity were responsible for decreased light sensitivity^17^, but whether RA-dependent remodeling corrupts image-forming vision has been unknown. Here, we found that in the primary visual cortex, disulfiram and BMS 493 treatment both increased the prevalence of orientation-selective neurons and sharpened their tuning. The recovery of fundamental response properties in the primary visual cortex indicates that elevation of RA signaling in the retina has detrimental downstream effects throughout the visual system.

### Relevance for human vision disorders

Indirect evidence strongly suggests that RA-induced retinal hyperactivity contributes to human vision impairment^24^. Direct evidence, however, is difficult to obtain because methods for detecting hyperactivity are invasive and therefore inappropriate for humans. In animal models of RP, hyperactivity can be detected directly with electrophysiological recordings from isolated retina, but RP is a rare disease, greatly restricting the availability of post-mortem human retinal samples for electrophysiological analysis. Non-invasive methods for recording activity in the human visual system include field potential recordings from retina (ERG) or brain (EEG), and radiological methods such as fMRI or PET scans. However, these only detect stimulus-evoked responses, providing no reliable measure of background activity.

Likewise, directly measuring heightened RA is fraught with difficulties. RAR binds to all- *trans* RA with a dissociation constant in the subnanomolar range (Kd = 0.4 nM^33^), suggesting that the ambient concentration of RA is very low. In addition, RA is labile owing to rapid enzymatic degradation by enzymes of cytochrome P450 family 26 (Cyp26)^52^. The combination of low concentration and short half-life make direct measurement of RA practically impossible, particularly in subregions of the retina (e.g., AMD and geographic atrophy). An alternative is indirect detection, for example with a RAR reporter gene incorporating the RARE sequence to drive fluorescent protein expression. However, delivery of a non-therapeutic reporter gene is considered too invasive and risky for implementation in humans, at least at present. Unfortunately, there are no suitable non-human primate models of inherited photoreceptor degeneration or AMD, limiting direct validation of heightened RA signaling to other mammalian models.

These issues make our results with disulfiram particularly exciting and compelling. Because disulfiram is FDA-approved with safety that has been established over decades^36^, it should face low regulatory hurdles for trials in humans with RP or other disorders, such as dry AMD. Our results on vision-impaired mice show that disulfiram improves behavioral contrast-sensitivity, sharpens cortical neuron representations of spatial orientation, and increases the fidelity of responses to naturalistic scenes, all consistent with improved visual perception. Whether disulfiram will improve vision in humans remains to be seen, but the barriers to answering this question seem relatively low. If disulfiram shows efficacy, it could be administered orally, but local ocular delivery involving a new drug formulation might ultimately be more appropriate for avoiding the undesired consequences associated with alcohol consumption. Disulfiram non-selectively inhibits all ALDH isozymes, but new drug candidates selectively targeting RALDH isoforms in the retina^53^ might also help remove these concerns.

Our results show that inhibiting RAR also improves vision. We used BMS 493, a pan-RAR inverse agonist that acts on RAR co-repressor protein^33, 40^. BMS 493 is potent and effective, but there is a rich pharmacopoeia of other small molecule agents that also interact with the RA binding site on RAR, the co-repressor or the co-activator, acting as agonists, antagonists, or inverse agonists. There are also agents that inhibit RXR, a protein that forms an obligate dimer with RAR before binding to DNA^54^. Determining which agent will have the most appropriate pharmacological properties for therapeutic vision rescue requires further studies of efficacy, toxicity, pharmacokinetics, and pharmacodynamics.

RAR expression or activity can also be inhibited by gene therapy. We have shown that a dominant-negative mutant of the RARα isoform (RAR_DN_), delivered to RGCs with a targeted AAV vector, suppresses hyperactivity and augments light responses in rd10 mice^17^. AAV-mediated gene delivery has the advantage of allowing cell-type targeting, which may benefit from the discovery of gene regulatory elements that may be expressed preferentially retinal neurons most severely impacted by RA-induced remodeling. However, in non-human primates, AAV-mediated gene delivery is limited by uneven penetration of the virus across the inner limiting membrane, restricting viral transduction to a ring of RGCs surrounding the fovea^55^. This scenario will presumably apply to humans as well. To avoid viral delivery limitations, other genetic tools may be appropriate for suppressing RAR expression, including RNA-interfering molecules such as antisense oligonucleotides (ASOs) and small interfering RNAs (siRNAs). These molecules are much smaller than viruses, and after lipid modification they can cross the inner limiting membrane and inhibit gene expression across the extent of the retina^56, 57^. ASOs and siRNAs are in clinical trials for other eye disorders and advances in their design may generally benefit this strategy for suppressing RA signaling.

In principle, light responses could be unmasked, and vision improved by directly inhibiting the ion channels that underlie hyperactivity. Gap junction proteins^58^, P2X receptors^20^, and HCN channels^11^ have all been implicated in remodeling, making them potential drug targets for vision improvement. However, these channels are all normally expressed in the healthy retina, and it might be impossible to block them without affecting normal function. It is possible that RA is the common signal for upregulating these channels, in which case a single RA inhibitor might reduce their expression in a coordinated manner. RAR is largely inactive in adult RGCs^17^, so an RAR inhibitor might reverse hyperactivity in degenerated regions of the retina without affecting normal function in regions with healthy photoreceptors.

Outer segments of rods and cones completely disappear in no-light perception patients with end-stage RP. However, at least some patients retain cone cell bodies that continue to express cone-specific genes, and these cells remain synaptically connected to bipolar cells^59^. The hyperactivity of downstream RGCs may obscure small light responses produced by these remnant cones, raising the possibility that elevated RA might obscure perception. Thus, there is the possibility that RALDH or RAR inhibitors might awaken light perception in patients otherwise considered completely blind.

Since RAR is relatively inactive in healthy RGCs, RAR inhibitors should act preferentially on regions of the retina where degeneration has taken place, bypassing healthy retinal regions. RA-induced gene expression is not limited to inherited retinal degeneration; it also occurs in response to photoreceptor degeneration caused by photodamage^21^ and local physical detachment from the retinal pigment epithelium^23^. This suggests that corruption of retinal processing may apply to many disorders that cause loss of rods and cones, perhaps including AMD.

Rescue of visual function by RAR may extend useful vision for months or years, but there is no evidence that it will slow photoreceptor degeneration itself. The strategy of rescuing vision with RA inhibitors is distinct from the strategy of restoring vision with retinal prosthetics, optogenetic and opto-pharmacological tools, or cell-based therapies. Vision restoration therapies are aimed, at least for now, at the small fraction of patients with end-stage photoreceptor degeneration, but treatments targeting the RA pathway may be relevant to the much larger patient population with low vision. Moreover, reducing RGC hyperactivity with RA inhibitors might be beneficial even after all the photoreceptors have degenerated and light perception is absent. Responses evoked by opto-electric^60, 61^, optogenetic^8, 62^, or optopharmacological^63^ stimulation of the degenerated retina are superimposed on the heightened background activity of RGCs, curtailing the encoding of visual images. The combination of a light-sensitive actuator with an RAR inhibitor could have a synergistic effect, boosting neural signals to more effectively restore visual function to no light perception patients.

## Materials and Methods

### Animals

C57/Black (‘wild-type’, WT) and rd10 (*Pde6β^-/-^* mutant) mice were purchased from Jackson Labs (strain #000664 and #004297, accordingly). All mice were kept in a 12:12 light/dark cycle room, except for experiments including dark rearing (see below). All animal procedures were approved by the Institutional Animal Care and User Committee at UC Berkeley and UC Santa Barbara.

### Chemicals and solutions

Dissections and ex-vivo retinal assays (MEA and imaging) were performed in artificial cerebrospinal fluid (ACSF) containing (in mM) 119 NaCl, 2.5 KCl, 1 KH_2_PO_4_, 1.3 MgCl_2_, 2.5 CaCl_2_, 26.2 NaHCO_3_, and 20 D-glucose. Disulfiram (tetraethylthiuram disulfide, C_10_H_20_N_2_S_4_) was purchased from Sigma (#86720) and formulated in mouse chow by Dyets Inc. (PA, USA), at a final concentration of 2 mg/Kg. Food was refreshed every 4-5 days during treatment. BSM 493 was purchased from Tocris (#3509) and stored at - 20℃ in a final concentration of 5 μM in PBS.

### Viruses

#### Retinoic Acid Receptor (RAR) reporter

To detect RAR-induced transcription, we used an AAV vector (Vigene Biosci., Maryland, USA) which included a cytomegalovirus promoter (CMV) upstream to the coding sequence for the red fluorescent protein (RFP, ‘mStrawberry’), followed by poly-A tail and a stop sequence^17, 23^. A fragment containing three repetitions of the retinoic acid response element (RARE) sequence followed by the weak promoter SV40 was sub-cloned from pGL3-RARE-luciferase (Addgene Plasmid #13458), a kind gift of the Underhill Lab^64^. Finally, a green fluorescent protein (GFP) sequence was sub-cloned downstream to SV40, for a final construct of AAV2-CMV-RFP-stop-RARE(x3)-SV40-GFP. The presence of inverted terminal repeat sequences was confirmed by enzymatic digestion.

#### Genetically-encoded calcium indicator

To measure activity-dependent increases in intracellular Ca^2+^ resulting from spike activity, we used a commercially available AAV vector encoding the Ca^2+^ indicator GCaMP6s under the control of the CaMKIIa promoter (Addgene #107790, AAV9- CamKII-GCaMP6s-WPRE-SV40), which is expressed in excitatory neurons.

### Intravitreal injections

Adult WT and rd10 mice (>P21) were intravitreally injected with drugs or virus. Before injection, animals were anesthetized with isoflurane (2%) and their pupils were dilated with tropicamide (1%) and phenylephrine (2.5%). Proparacaine (0.5%) was used as a topical analgesic. Genteal was applied under a glass coverslip to keep the cornea lubricated. An incision was made through the sclera below the ora serrata with a 30G needle. Solutions were injected into the vitreous with a blunt-ended 33G Hamilton syringe or a 25 - 30μm tip diameter glass pipette beveled at a 45° angle. After injection, the antibiotic tobramycin (0.3%) was applied to the eye. Injections of RAR-reporter AAV2 were binocular, at a maximal volume of 1.5 μl, and using a titer of >10^14^ particles/μl. Injections of vehicle (PBS x1) and BMS 493 (5 μM) were binocular or monocular (depending on the experiment), using a final volume of 1.0 μL. Final BMS 493 concentration was 500 nM, assuming a 1:10 dilution in the vitreous volume of the mouse (∼10 μL).

### Retinal dissections

All dissections were performed in ACSF continuously gassed with 95% O_2_/5% CO_2_. Mice were euthanized using saturating levels of isoflurane gas followed by cervical dislocation. Mice were then enucleated, the cornea was excised from the eye and the lens removed. For live retinal imaging and MEA recordings, retinas were detached from the retinal pigment epithelium and cut in quarters. Each retinal piece was mounted on nitrocellulose paper, RGCs facing up (imaging), or placed on top of the recording electrodes, RGCs facing down (MEA). Retinal pieces were kept at 34-35℃.

### Cortical surgery

All surgeries were conducted with the mouse under isoflurane anesthesia (3.5% induction, 1.5-2.5% maintenance). Prior to incision, the scalp was infiltrated with lidocaine (5 mg/kg, subcutaneous) for analgesia. Meloxicam (2 mg/kg, subcutaneous) was administered preoperatively to reduce inflammation. Once anesthetized, the scalp overlying the dorsal skull was sanitized and removed. The periosteum was removed with a scalpel and the skull was abraded with a drill burr to improve adhesion of dental acrylic.

For mice in the disulfiram experiments, a single 4 mm craniotomy was made over the left visual cortex. For mice in the BMS 493 experiments, a 4 mm craniotomy was made over each bilateral visual cortex (centered at 4.0 mm posterior, ±2.5 mm lateral to Bregma on each side), leaving the dura intact. We used a motorized microinjector (Stoelting, 53311) to deliver 0.75 μl of virus AAV9-CamKII-GCaMP6s (titer = 1-3×10^13^) to 3-4 injection sites within area V1 of the cerebral cortex. A cranial window was implanted over the craniotomy and sealed, first with silicon elastomer (Kwik-Sil, World Precision Instruments) and then with dental acrylic (C&B-Metabond, Parkell) mixed with black ink to reduce light transmission. The cranial windows were made of two rounded pieces of cover glass (Warner Instruments) bonded with a UV-cured optical adhesive (Norland, NOA61). The bottom cover glass (4 mm) fit tightly inside the craniotomy while the top cover glass (5 mm) was bonded to the skull with the dental acrylic. A custom-designed stainless-steel head plate (eMachineShop.com) was then affixed using dental acrylic. After surgery, mice were administered carprofen (5-10 mg/kg, oral) every day for 3 days to reduce inflammation. The full specifications and designs for head plate and head fixation hardware can be found on our institutional lab website (https://labs.mcdb.ucsb.edu/goard/michael/content/resources).

### RAR Reporter Imaging Assay

Live retinal pieces mounted on nitrocellulose paper were maintained under oxygenated ACSF perfusion at 34℃. A spinning-disk confocal microscope (Olympus IX-50) with a 40x water-submersible objective was used for fluorescence imaging to detect red or green fluorescence owing to RFP or GFP expression. 1.5 μm-thick optical sections of the ganglion cell layer of the retina were compiled to generate Z-stacks. Z-stacks were flattened and analyzed with ImageJ (NIH). Two to 3 fields of view were analyzed for each retinal piece and individual regions of interest (ROIs) were drawn around every visible cell body, enabling measurement of mean grey value (MGV) for both RFP and GFP fluorescence. The GFP/RFP ratio was calculated for each ROI and averaged across retinal pieces (individual data points) and mice (mean value).

### MEA

Retinas were dissected and maintained in ACSF as described previously. Individual pieces of retina were placed ganglion cell layer down onto an array with 60 electrodes spaced 200 μm apart (1060-2-BC, Multi-Channel Systems). After mounting, each retinal piece was dark-adapted for 30 minutes under constant perfusion of 34°C oxygenated ACSF. Extracellular signals were digitized at 20 kHz and passed through a 200 Hz high-pass 2nd order Butterworth recursive filter. Spikes were extracted using a threshold voltage of 4 SD from the median background signal of each channel. Spikes were then aligned and clustered primarily in 3D principal component space using T-Distribution Expectation-Maximization (Offline Sorter, Plexon). Inclusion criteria for units included distinct depolarization and hyperpolarization phases, inter-spike interval histograms with peak values, and at least 50 contributing spikes. Exclusion criteria included multiple peaks, high noise, and low amplitude in channels with more than 3 detected units.

### Electroretinogram

Photopic ERG responses were recorded in adult mice with the HMsERG LAB System (OcuScience), under isoflurane anesthesia (3.5% induction, 1.5 - 2.5% maintenance) at a stable rectal temperature of >35.8℃. The ground electrode was placed on the back above the tail, while reference electrodes were placed in the chick under the cheek (all subdermal). Mouse pupils were dilated with tropicamide (1%) and phenylephrine (2.5%). Proparacaine (0.5%) was used as a topical analgesic. To create and maintain contact between the lens containing the recording electrode and the cornea, a small drop of hypromellose was placed on the eye and the lens was gently pressed against it. After recording, Genteal was applied to both eyes and mice were allowed to recover from anesthesia.

Photopic stimulation was performed inside a mini-ganzfeld stimulator including 10 minutes of light adaptation to background light at 300 cd.s/m^2^, followed by 32 flashes (frequency = 2.0 Hz, duration = 20 msec) for 8 different light intensities ranging from 0.1 - 25 cd.s/m^2^. Responses were recorded over a period of 360 msec after each flash. Data were filtered at 50 KHz. The amplitude of the b-wave was measured from the peak of the first inward voltage deflection (a-wave) to the peak of the first outward voltage deflection and averaged across 32 flashes per intensity.

### Visual Behavioral Assay

#### Cage design and manipulation

The automated behavioral cages were based on a design developed by Ed Pugh^44^, and custom-built by Lafayette Instrument (IN, USA). The cage includes three main components: i) a unidirectional running wheel, orienting the mouse towards a computer display and a revolutions counter, ii) a nose poke area with an infrared sensor, connected to a peristaltic pump triggering delivery of a water droplet reward, and iii) a Raspberry Pi mini-computer and 1366 x 768 pixels LCD screens. Screens were calibrated to have the same brightness. Custom software linked image display with wheel running and reward availability. Cages were used in a dark room and experiments were conducted throughout the night. The involvement of the animal handlers was minimal (introduction and retrieval), while protocol implementation and data acquisition were fully automated.

#### Behavioral experiment design and data analysis

Adult (P>35), male and female WT and rd10 mice, in littermate tandems, were used for these experiments. The cohort of rd10 mice used in disulfiram experiments (**Fig. 4**) were reared in 14:10 light-dark conditions. The cohort of rd10 mice used in BMS 493 experiments (**Fig. 5**) were dark reared from P<10 to P55-65 (depending on training and testing), and under 14:10 light-dark conditions afterwards. Mice were water-deprived for >9 hours ahead of each experiment. Mice learned the task through operant conditioning. Training and testing sessions were conducted for 12 hours (6 PM to 6AM). During the habituation protocol, the visual stimulus (a full-contrast grating with a 0.51 CPD frequency) associated with reward availability is continuously, and the mouse can poke and receive 10% sucrose water *ad libitum*. During training, running on the wheel gates the visual stimulus, displayed on the screen. This is paired with unlocking of the nose poke, enabling the mouse to obtain the reward. The reward availability period, and the duration of pump activation are shortened during training sessions to increase response stringency. During testing, stimuli have the same total luminance, but the contrast is randomly chosen from 9 different possible steps, including 0, 1, 2, 4, 8, 16, 32, 64 and 100%.

In all sessions, latency was defined as the elapsed time between onset of the visual stimulus to nose poke. The mean latency to the 100% contrast stimulus during the first contrast-sensitivity test was used as a reference to assess performance in response to all other contrasts and for all subsequent tests. A successful response was defined as one with mean latency falling within the ±2SD of the latency during the reference test. Exclusion criteria during training: if during session 3, success rate was <80% and/or mean latency was >4 sec. Exclusion criteria during testing: if SD >40% of the mean (SD/mean>0.4, see **Table S5 and S6**). Initial analysis was performed with custom software to extract the data from text files and arrange responses by contrast steps in datasheets. Analysis of success rate was performed manually.

### Cortical Imaging

#### Visual stimuli

All visual stimuli were generated with a Windows PC using MATLAB and the Psychophysics toolbox (Brainard, 1997). Visual stimuli were presented on two LCD monitors that can display visual images to either eye independently. Each monitor (17.5 × 13 cm, 800 × 600 pixels, 60 Hz refresh rate) was positioned symmetrically 5 cm from each eye at a 30° angle right of the midline, spanning 120° (azimuth) by 100° (elevation) of visual space. The monitors were located 3 cm above 0° elevation and tilted 20° downward. A nonreflective drape was placed over the inactive monitor to reduce reflections from the active monitor.

Orientation tuning was measured with drifting sine wave gratings (spatial frequency 0.05 cycles/deg; temporal frequency: 2Hz) presented in one of 12 directions, spanning from 0 degrees to 330 degrees in 30 degree increments. For a single repeat, each grating was presented once for 2 seconds with a luminance matched 4 second inter-trial gray screen between presentations. This was repeated for 8 repetitions per session.

Natural scenes stimuli consisted of 900 frames from *Touch of Evil* (Orson Wells, Universal Pictures, 1958) presented at 30 frames per second, leading to a 30 second presentation time per repeat with a 5 second inter-trial gray screen. The clip consists of a single continuous scene with no cuts, as has been previously described and is commonly used as a visual stimulus (https://observatory.brain-map.org/visualcoding/stimulus/natural_movies). Each presentation was repeated 30 times, with a 5 second inter-trial gray screen.

#### Two-photon imaging

After >2 weeks recovery from surgery, GCaMP6s fluorescence was imaged using a Prairie Investigator two-photon microscopy system with a resonant galvo-scanning module (Bruker). For fluorescence excitation, we used a Ti:Sapphire laser (Mai-Tai eHP, Newport) with dispersion compensation (Deep See, Newport) tuned to λ = 920 nm. For collection, we used GaAsP photomultiplier tubes (Hamamatsu). To achieve a wide field of view, we used a 16X/0.8 NA microscope objective (Nikon) at 1X (850 × 850 μm) or 2X (425 × 425 μm) magnification. Laser power ranged from 40 to 75 mW at the sample depending on GCaMP6s expression levels. Photobleaching was minimal (<1%/min) for all laser powers used. A custom stainless-steel light blocker (eMachineShop.com) was mounted to the head plate and interlocked with a tube around the objective to prevent light from the visual stimulus monitor from reaching the PMTs. During imaging experiments, the polypropylene tube supporting the mouse was suspended from the behavior platform with high tension springs (Small Parts) to reduce movement artifacts.

#### Two-photon post-processing

Images were acquired using PrairieView acquisition software and converted into TIF files. All subsequent analyses were performed in MATLAB (Mathworks) using custom code (https://labs.mcdb.ucsb.edu/goard/michael/content/resources). First, images were corrected for X–Y movement by registration to a reference image (the pixel-wise mean of all frames) using 2-dimensional cross correlation

To identify responsive neural somata, a pixel-wise activity map was calculated using a modified kurtosis measure. Neuron cell bodies were identified using local adaptive threshold and iterative segmentation. Automatically defined ROIs were then manually checked for proper segmentation in a graphical user interface (allowing comparison to raw fluorescence and activity map images). To ensure that the response of individual neurons was not due to local neuropil contamination of somatic signals, a corrected fluorescence measure was estimated according to:

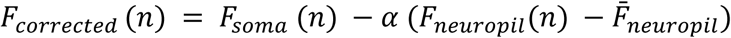

where *F*_neuropil_ was defined as the fluorescence in the region <30 μm from the ROI border (excluding other ROIs) for frame *n* and *α* was chosen from [0 1] to minimize the Pearson’s correlation coefficient between *F*_corrected_ and *F*_neuropil_. The Δ*F*/*F* for each neuron was then calculated as:

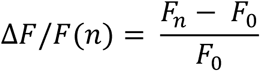

where *F_n_* is the corrected fluorescence (*F*_corrected_) for frame *n* and *F*_0_ defined as the first mode of the corrected fluorescence density distribution across the entire time series.

### Analysis of two-photon imaging data

#### Blinding to experimental condition

For disulfiram experiments, disulfiram-containing chow or control chow of the same composition (Dyets, Inc) were given neutral codes by an investigator not involved in the study and administered to the mice with the experimenter blind to experimental condition. For BMS-493 experiments, vials of drug and vehicle were given neutral codes and each solution was used for one eye. In both cases, the experimental condition was revealed only after the primary analysis was complete.

#### Orientation tuning

Neural responses to the orientation tuning stimulus were first separated into trials, each containing the response of the neuron across all tested queried orientations. For each neuron, we averaged the baseline-subtracted responses to each orientation, creating an orientation tuning curve for each trial. To calculate the orientation selectivity index (OSI) in a cross-validated manner, we first separated the orientation tuning curves into even and odd trials. We then aligned the even trials using the maximal response of the averaged odd trial for each neuron, and vice versa, resulting in aligned responses. We then calculated the OSI from the averaged tuning curves for each neuron using the following equation:

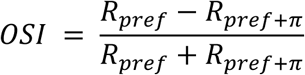

where *R_pref_* is the neuron’s average response at its preferred orientation, defined by cross-validation on a different set of trials using the above procedure. Aligning the orientation tuning curves of the neurons using cross-validation provides a more accurate measurement of the orientation tuning of the neuron, as it prevents non-selective neurons from having high OSI values due to spurious neural activity.

#### Naturalistic movies reliability

Neural responses to the naturalistic movies were first separated into trials, with each trial containing the full response of the neuron to the entire presented movie. The reliability of each neuron to the naturalistic movie was calculated as follows:

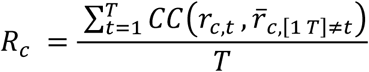

where *R* is the reliability for cell *c,t* is the trial number from [1 *T*], *CC* is the Pearson correlation coefficient, *r_c,t_* is the response of cell *c* on trial *t* and 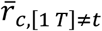 is the average response of cell *c* on all trials excluding trial *t*.

#### Naturalistic movies decoding analyses

To decode naturalistic movie responses from the population data, we first randomly selected neurons of a given pool size (pool sizes: 2, 4, 8, 16, 32, 64, 128, 256, or 512 neurons) from across all recordings. The neural responses to natural movies within that pool were then divided into even and odd trials. The average population activity across even trials was used to calculate a “template” population vector for each frame of the movie. We then estimated the movie frame (*F_Decoded_*) from the population activity of each actual frame (*F_Actual_*). To accomplish this, we calculated the population vector from the odd trials during *F_Actual_* and compared to the “template” population vectors (even trials) for all of the frames. The frame with the smallest Euclidean distance between population vectors was chosen as the decoded frame (*F_Decoded_*). This process was repeated for each frame (*F_Actual_*) of the movie. For each pool size of neurons used, the entire procedure was iterated 1000 times, picking new neurons for each iteration. This resulted in a confusion matrix that describes the similarity of neural activity patterns for each frame between non-overlapping trials. To assess decoder performance, we measured the percentage of decoded frames that fell within 10 frames of the actual frame (chance level = 7%).

### Statistical Analysis

The specific statistical test employed in each dataset is specified in its corresponding figure legend and also detailed in **Tables S1,3,5,6.** Comparison between groups used non-parametric tests (Mann-Whitney U test for independent samples or Wilcoxon sign test for paired data), unless the data passed the normality test (Shapiro-Wilk) and could be analyzed with parametric tests. Significance between cumulative probabilities was tested using the Kolmogorov–Smirnov test. Two samples were considered significantly different when the p value was <0.05.

#### Linear Mixed Effects Modeling

Because of the highly nested nature of physiological recordings, we employed linear mixed effects models (LMEMs) to account for the individual differences between mice in our statistical analyses. The formula for the LMEM is as follows:

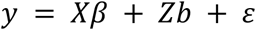

where *y* is the response vector, *X* is the fixed-effects design matrix (denoting treatment condition), *β* is the fixed effects vector, *Z* is the random-effects design matrix (denoting different mice), *b* is the random effects vector, and ε is the observation error vector.

After fitting the LMEMs for each experiment, we performed a one-sample t-test on the main effect of “treatment condition” to gauge the significance of our experimental treatments.

## Supporting information

Legends for Supp Figures and Tables

## Author Contribution

M.T. experimental design, data collection and analysis for RAR-reporter assay, MEA recordings, visual behavior assay; manuscript writing and editing.

K.S. experimental design, data collection and analysis for cortical calcium imaging experiments, manuscript writing and editing.

D.F. mouse husbandry, data collection and analysis for MEA recordings, data collection for visual behavior assay.

B.S.experimental design for visual behavior assay.

A.M. data analysis for visual behavior assay.

M.J.G. experimental design, manuscript writing and editing, funding.

R.H.K.experimental design, manuscript writing and editing, funding.

## Acknowledgements

We thank Mei Li and Hong Ma (UC Berkeley) for viral production and intravitreal injections, respectively.

