## Supplementary figures and images for "Inhibiting retinoic acid mitigates vision loss in a mouse model of retinal degeneration"

### Figure S1

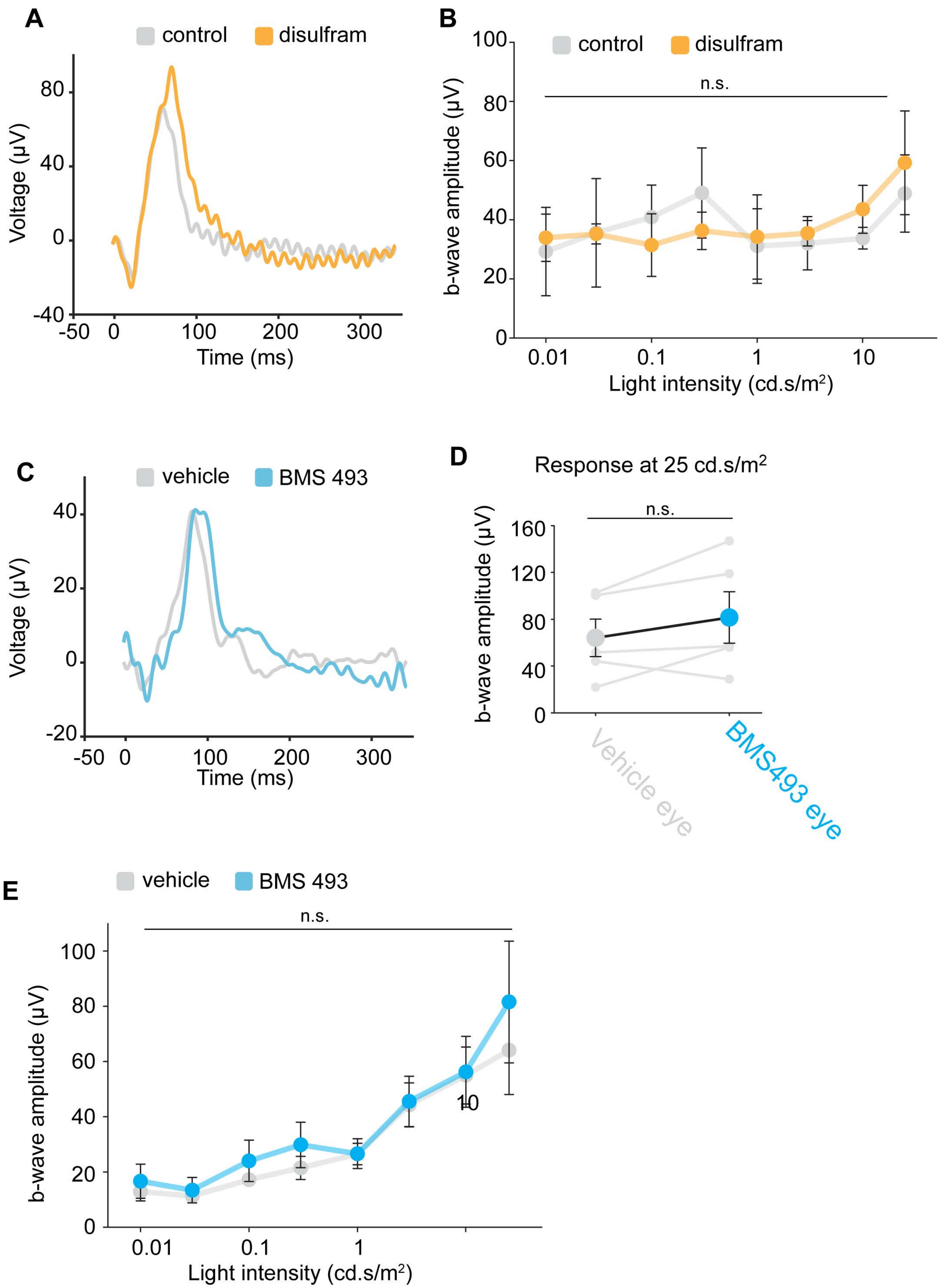

### Figure S2

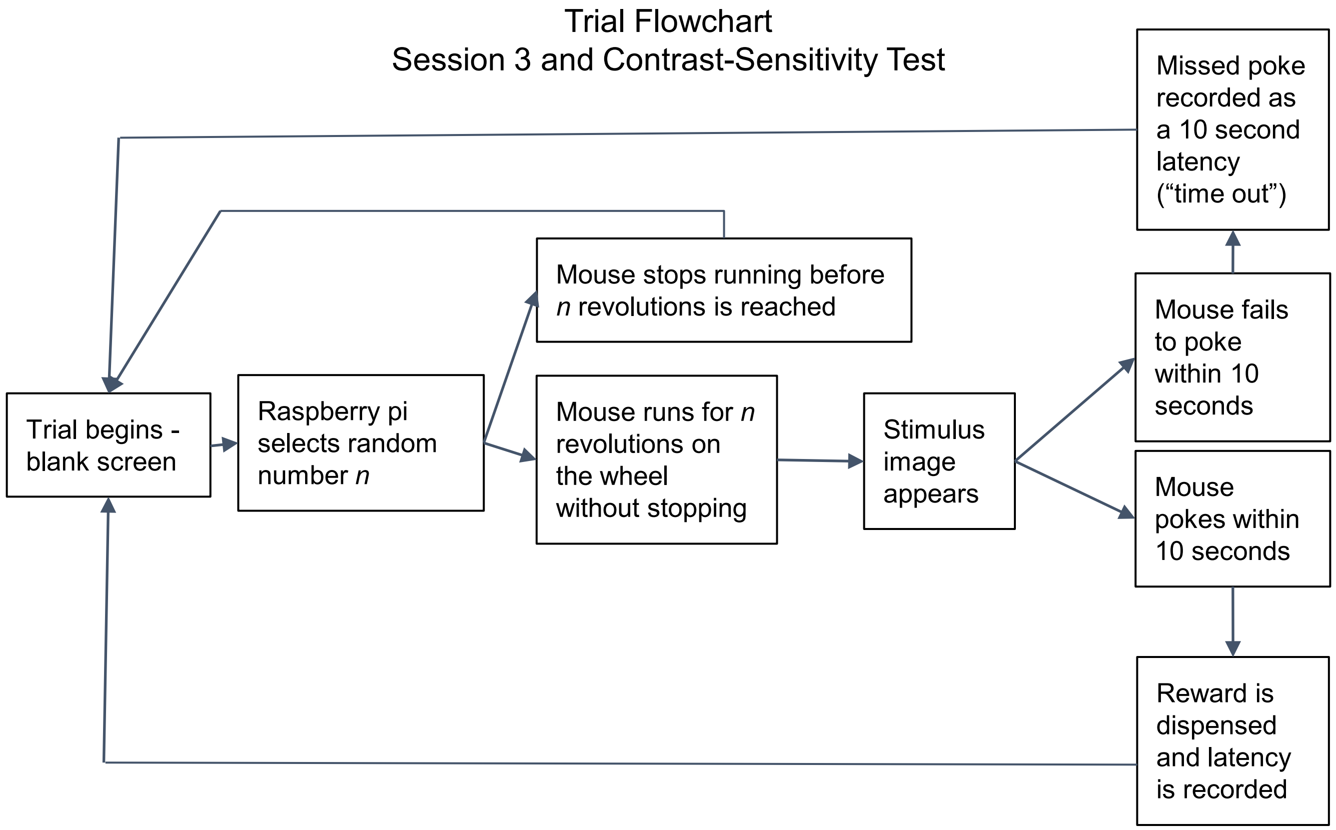

### Figure S3

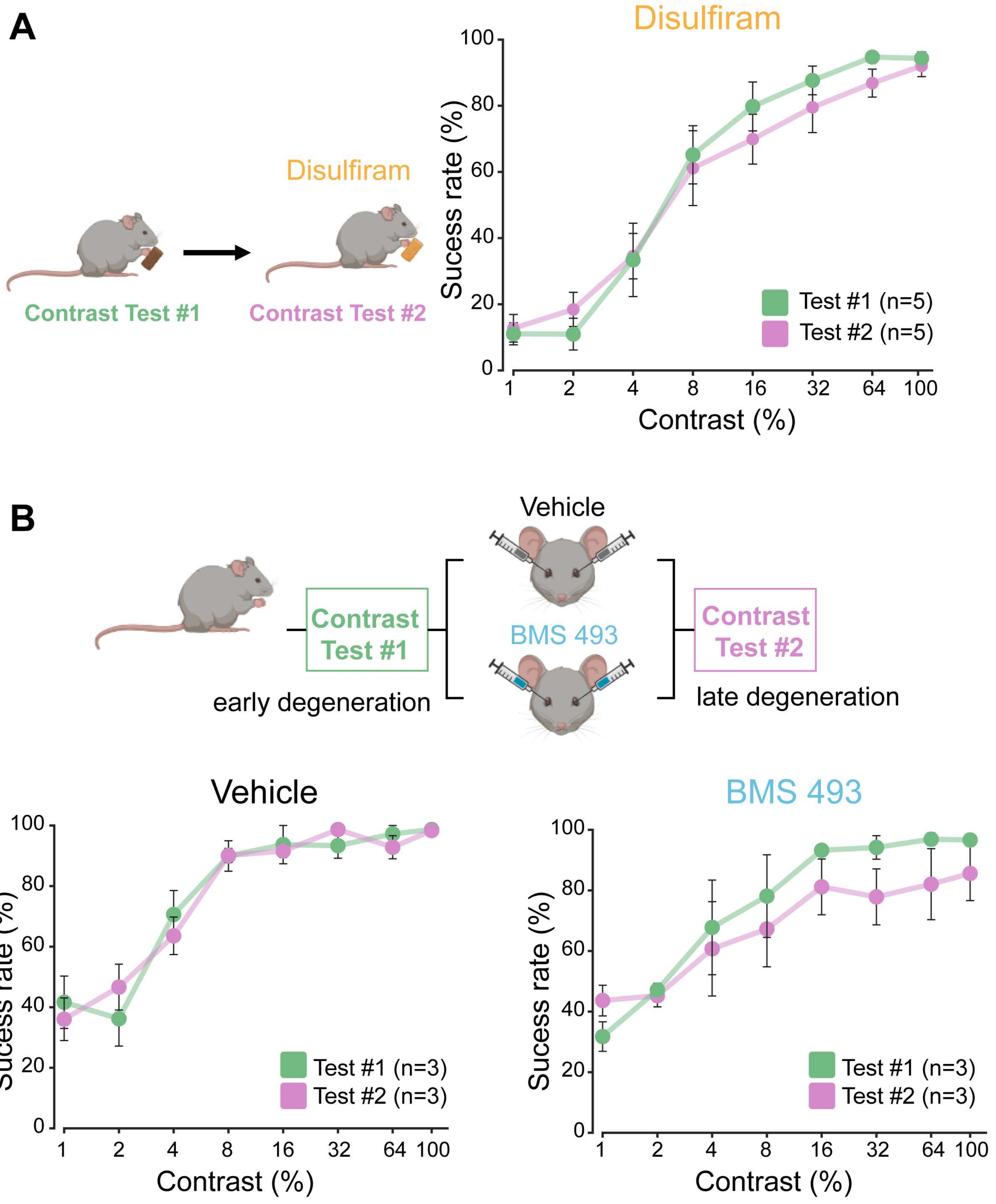
