## Supplementary material for "Inhibiting retinoic acid mitigates vision loss in a mouse model of retinal degeneration": Legends for Supp Figures and Tables

**Supplemental Material**

Legends to Supplemental Figures

**Supplemental Figure 1: The ERG b-wave is unchanged by treatment with disulfiram or BMS 493.
A)** Electroretinogram (ERG) recordings of P80-90 rd10 mice treated for 40 days with disulfiram (orange) vs control (grey). Traces are the mean response to 32 flashes of light (25 cd.s/m^2^ light for 20 msec) presented at a frequency of 2 Hz. **B)** Intensity-response curves showing no significant effect of disulfiram. The amplitude of the b-wave was measured from the peak of the first inward voltage deflection (a-wave) to the peak of the first outward voltage deflection. Recordings were obtained from 3 control mice (6 eyes) and 4 disulfiram mice (8 eyes) and are shown as mean ± SEM. n.s. - non-significant difference, p>0.05, 2-tailed t-test (**Table S3**). **C)** ERG recordings 4-6 days after injecting vehicle (PBS x1, grey) into one eye and BMS 493 (blue) into the contralateral eye. **D)** Responses from BMS 493-injected and vehicle-injected eyes in all 5 mice tested. **E)** Intensity-response curves showing no significant effect of BMS 493. Recordings were obtained from 5 mice (10 eyes), one eye injected with BMS 493 and the contralateral eye with vehicle. Injections were randomized across eyes and mice. n.s. - non-significant difference, p>0.05, 2-tailed paired t-test (**Table S3**).

**Supplemental Figure 2: Parameters defining a single trial in a trained mouse.**

During session 3 and contrast sensitivity test (**Fig. 3B-C**), the stimulus is displayed for a maximum of 10 seconds after it is triggered by wheel revolutions. Each trial begins with a blank screen displaying a 0% contrast version of the stimulus image (‘blank screen’). The computer (i.e.: raspberry pi) chooses a random number of wheel revolutions (see **Table S4**) necessary to display the stimulus image. Stimulus image is always 100% contrast in sessions 1-3, or 1 out of 9 different contrasts, randomly chosen, during contrast sensitivity test. The response of the mouse (i.e. nose poke) is registered, and the delivery of reward is activated if the mouse pokes before “time out” (10 seconds for training session 3 and contrast sensitivity test, see **Table S4**).

**Supplemental Figure 3**. Testing disulfiram and BMS 493 in wild-type mice with normal vision.

**A**) Left: experimental design. C57/Black male mice were trained and tested at ~P60, using the same behavioral paradigm as shown in **Fig. 3A-C**. Mice were fed a regular diet without disulfiram and tested for contrast sensitivity (test #1), after which their food was switched to that containing disulfiram (2 mg/Kg) for 40 days. At ~P100 they were tested for the second time (test #2). Right: success rate for each contrast tested before (green) and after (pink) disulfiram treatment. Values are shown as mean ± SEM, p>0.05 in all contrasts, Mann-Whitney test. **B**) Similar to **A**, above. Six C57/Black male mice were trained and tested at P60, and then injected intravitreally in both eyes with 1 μl of vehicle (PBS x1) or BMS 493 (5 μM). At 4-6 days post-injection, they were tested for a second time. The success rate in test #1 was not significantly different from the success rate during test #2 in either of the two groups. Values are shown as mean ± SEM, p>0.05 in all contrasts, Mann-Whitney test.

Legends to Supplemental Tables

**Supplemental Table 1. Data and statistical analysis corresponding to Figure 1.**

In disulfiram vs control experiments (**Fig. 1E**), treatment was systemic and both eyes were dissected together. Each retina was cut into 3-4 pieces and 3-6 retinal samples were analyzed.

In BMS 493 vs vehicle experiments (**Fig. 1H**), in each individual mouse one eye was injected with the drug and the second with the vehicle control (see Materials and Methods). Each eye was dissected separately and cut into 4 pieces, 3-4 retinal samples were imaged and analyzed for each eye. Data obtained using ex-vivo fluorescent imaging in ACSF (see **Materials and Methods**).

**Supplemental Table 2. Data and statistical analysis corresponding to Figure 2.**

Individual data points for mean firing frequency (**Fig. 2C**), and cumulative probability plot (**Fig. 2D**) in control and disulfiram-treated mice, as well as in vehicle and BMS 493-injected mice (**Fig. 2G,H**). In addition the table provides the data for the total and relative number of active units in retinas in all four conditions. Data obtained using ex-vivo MEA recordings in ACSF (see **Materials and Methods**).

**Supplemental Table 3. Data and statistical analysis corresponding to Supplemental Figure 1.**

Individual data points and statistical analysis for the b-wave’s amplitude (in μV), used for the plots shown in **Fig. S1B,D,E**). Data obtained using in-vivo ERG recordings of anesthetized mice (see **Materials and Methods**).

**Supplemental Table 4. Parameters of training and testing protocols in the behavioral paradigm (corresponds to Figure 3).**

Detailed description of the parameters employed during each protocol for habituation, training and testing, in the mouse behavioral paradigm (**Fig. 3B,C**). Each protocol lasted 12 hours from 6 PM to 6 AM. Protocols were encoded in python and manually loaded onto each raspberry pie using signature dongles, each corresponding to a specific cage.

**Supplemental Table 5. Data for contrast-sensitivity test in rd10 mice in control vs disulfiram conditions (corresponds to Figure 4).**

Top: Data and statistical analysis of success rates in rd10 mice during test #1 and test #2, in control and in disulfiram-treated mice (**Fig. 4D,E**). Bottom: data on the inclusion and exclusion of mice in the behavioral paradigm experiment for Figure 4.

**Supplemental Table 6. Data for contrast-sensitivity test in rd10 mice in vehicle vs BMS 493 conditions (corresponds to Figure 5).**

Top: Data and statistical analysis of success rates in rd10 mice during test #1 and test #2, in vehicle and in BMS 493-injected mice (**Fig. 5D,E**). Bottom: data on the inclusion and exclusion of mice in the behavioral paradigm experiment for Figure 5.
